# The role of m6A-RNA methylation in stress response regulation

**DOI:** 10.1101/200402

**Authors:** Mareen Engel, Simone Röh, Carola Eggert, Paul M. Kaplick, Lisa Tietze, Janine Arloth, Peter Weber, Monika Rex-Haffner, Mira Jakovcevski, Manfred Uhr, Matthias Eder, Carsten T. Wotjak, Mathias V. Schmidt, Jan M. Deussing, Elisabeth B. Binder, Alon Chen

## Abstract

N6-Methyladenosine (m6A) is an abundant internal RNA modification that regulates transcript processing and translation. The regulation of brain m6A by stressful stimuli in vivo and its role in the stress response are currently unknown.

Here, we provide a detailed analysis of the stress-epitranscriptome using m6A-Seq, global and gene-specific m6A measurements. We show that stress exposure and glucocorticoids alter m6A and its regulatory network in a region- and time-specific manner. We demonstrate that depletion of the methyltransferase Mettl3 and the demethylase Fto in adult neurons increases fear memory, and alters the transcriptome response to fear as well as synaptic plasticity. Finally, we report that regulation of m6A is impaired in major depressive disorder patients following glucocorticoid receptor activation.

Our findings indicate that brain m6A represents a novel layer of complexity in gene expression regulation after stress and that dysregulation of the m6A-response may contribute to the pathophysiology of stress-related psychiatric disorders.

**Highlights:**

- m^6^A RNA methylation in adult mouse brain is regulated by stress
- Brain m^6^A levels are temporally and spatially regulated by stress
- Mettl3 and Fto-KO alter fear memory, transcriptome response and synaptic plasticity
- The m^6^A-glucocorticoid-response is impaired in major depressive disorder patients

**eTOC blurb:** Engel et al. demonstrate a brain-area-specific and time-dependent role for the mRNA modification, m^6^A, in stress-response regulation. Manipulating m^6^A-enzymes alters fear-memory, transcriptome-response and synaptic-plasticity. Altered m^6^A dynamics in depressed patients suggest an involvement of m6A-modifications in stress-related psychiatric disorders.

## Introduction

Regulation of gene expression in response to stressful stimuli under healthy or pathological conditions involves epigenetic mechanisms such as DNA methylation and chromatin modifications (de Kloet et al., 2005; McEwen et al., 2015). Elucidating the underlying molecular processes that regulate the fine-tuned transcriptional response to stress is essential for understanding stress vulnerability and the development of stress-related psychiatric disorders such as depression and anxiety.

In analogy to DNA modifications, a diverse set of covalent modifications is present on RNA nucleotides, encoding the epitranscriptome and regulating gene expression post-transcriptionally. These modifications determine how much protein is translated from available mRNA and how non-coding transcripts function (Zhao et al., 2017). However, the role of this newly emerging layer of gene expression control in the central stress response and behaviour is not known yet. RNA modifications, next to epigenetic mechanisms, likely represent an yet undescribed level of transcriptional regulation highly relevant for psychiatry, and will contribute to our understanding of the combined effects of genetic and environmental factors in shaping disease risk (Klengel and Binder, 2015).

N^6^-methyladenosine (m^6^A) is the most abundant internal mRNA modification, which is present transcriptome-wide in at least one fourth of all RNAs, typically located in a consensus motif (DRACH/GGACU), and enriched near stop codons and in 5’UTRs (Dominissini et al., 2012; Linder et al., 2015; Meyer et al., 2012). Recent studies have identified mammalian m^6^A to be dynamically regulated, controlling stem cell proliferation and differentiation (Klungland et al., 2017), cellular heat-shock response (Zhou et al., 2015), DNA damage response (Xiang et al., 2017) and tumorigenesis (Cui et al., 2017). Brain RNA methylation is comparably high (Meyer et al., 2012) and increases during development (Meyer et al., 2012).

m^6^A is deposited co-transcriptionally (Ke et al., 2017; Slobodin et al., 2017) by a methyltransferase complex consisting of METTL3, METTL14 (Liu et al., 2014), WTAP (Ping et al., 2014), KIAA1429 (VIR) (Schwartz et al., 2014), and RBM15/RBM15B (Patil et al., 2016). In contrast, it can be removed by the demethylases FTO (Jia et al., 2011; Mauer et al., 2017) and ALKBH5 (Zheng et al., 2013). FTO further catalyses demethylation of N^6^,2′-O-dimethyladenosine (m^6^Am) with an *in vitro* preference for this substrate (Mauer et al., 2017). m^6^Am is found at the first nucleotide of certain RNAs adjacent to the 7-methylguanosine cap and promotes transcript stability (Mauer et al., 2017). The most commonly used anti-m^6^A antibody co-detects m^6^A and m^6^Am and therefore both m^6^A and m^6^Am are included in most of our measurements, henceforth both termed m^6^A without further specification. *Fto* has been associated with memory consolidation (Walters et al., 2017; Widagdo et al., 2016) and was implicated in regulation of dopaminergic brain networks (Hess et al., 2013). m^6^A enzymes are expressed at different levels in different cell-types and have distinct intracellular distributions and binding motifs and thus potentially affect different subsets of target RNAs. Cellular consequences of m^6^A modifications depend on the binding of m^6^A-reader proteins (such as YTH- and HNRNP-proteins) and include RNA maturation, potentially splicing (Alarcón et al., 2015; Ke et al., 2017; Liu et al., 2015; Xiao et al., 2016), alternative polyadenylation (Ke et al., 2015), RNA decay (Shi et al., 2017; Wang et al., 2014), as well as both promotion and inhibition of protein translation (Meyer et al., 2015; Shi et al., 2017; Wang et al., 2015; Zhou et al., 2015).

In this study, we aimed to elucidate the role of m^6^A in the context of the brain’s stress response. We delineated the effects of acute stress on m^6^A using global m^6^A measurements, m^6^A-Seq and absolute quantification of transcript-specific methylation levels. In addition, we explored the functional significance of m^6^A in the adult brain by examining conditional knockout mice for *Mettl3* and *Fto*. Finally, we investigated m^6^A regulation in blood samples of mice and humans to determine its potential as a peripheral indicator of the central response to stress and stress-linked psychiatric disorders.

## Results

### The stress-induced m6A epitranscriptome

To test whether acute stress alters m^6^A, we performed m^6^A-Seq (RNA-Seq after m^6^A-immunoprecipitation) on mouse cortex 4 h following 15 min of acute restraint stress exposure. To facilitate the detection of biologically significant changes, we applied a minimum cut-off on expression level of RNA RPKM>1 (Figure S1 A). We observed a total of 25,821 m^6^A-peaks consistent across 7 biological replicates per condition, mapping to 11,534 genes (Figure 1 A, Table S1). On average, each transcript had 2.24 peaks (Figure S1 B). Of these peaks, 55.4 % overlapped with previously reported m^6^A-peaks (RMBase (Sun et al., 2016)) but we found an additional 3,199 novel m^6^A-methylated genes. Although 1,810 m^6^A-peaks (mapped to 1644 genes) were differentially methylated after stress (Table S1), only 179 transcripts were found to be expressed in a stress-affected manner (FDR-corrected P-Value < 0.05, absolute log2 fold change > 0.1) (Figure 1 A). Interestingly, only 16% (28) of these genes were regulated both on the RNA and m^6^A-level. Both the absolute extents of m^6^A fold changes (Figure S1 A) as well as RNA fold changes were rather small, which may reflect the cellular heterogeneity of the input material.

**Figure 1.**
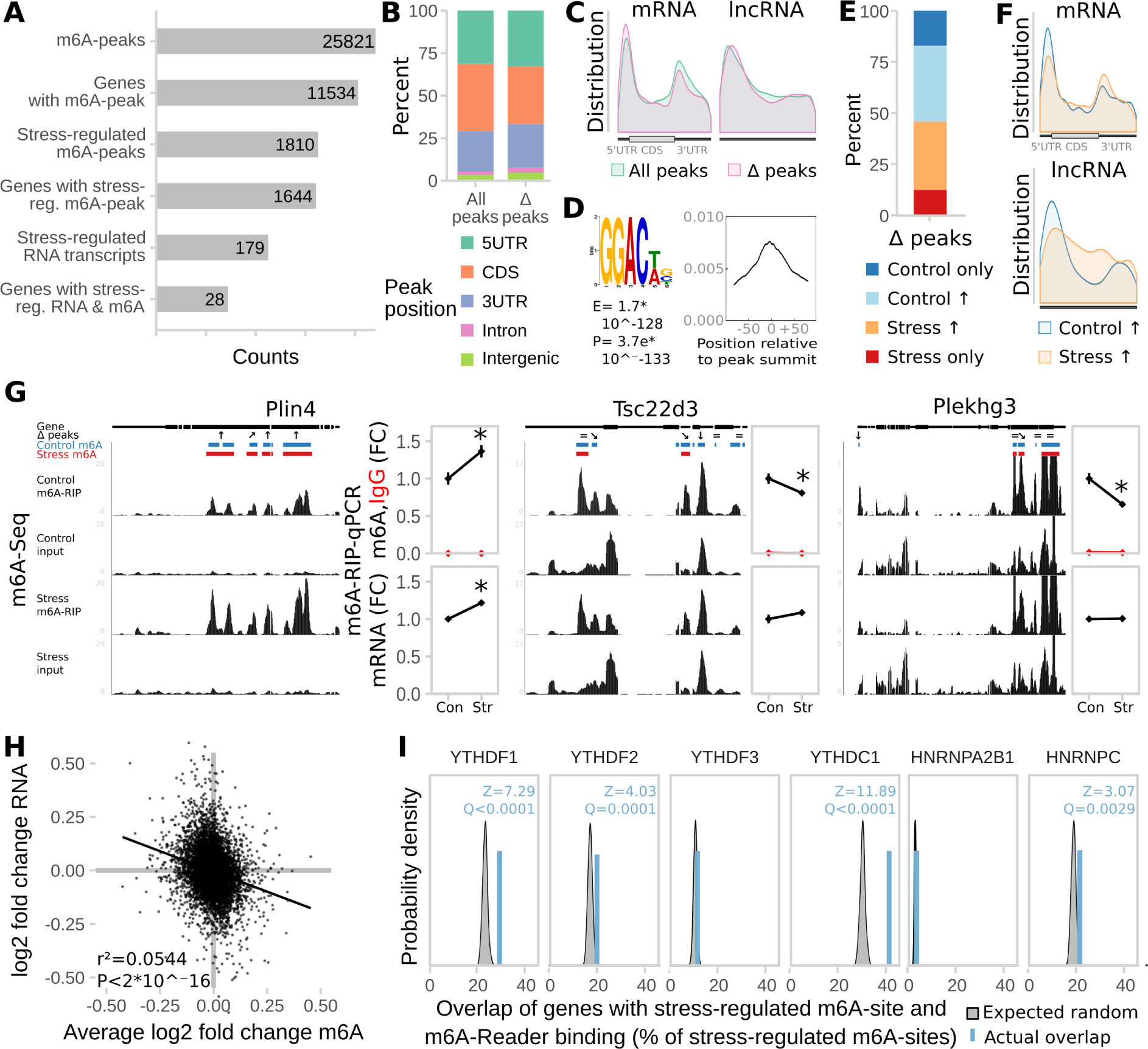
m6A-Seq reveals the transcriptome-wide map of stress-regulated m6A sites after acute restraint stress in the mouse cortex. **(A) The mouse cortical transcriptome is largely methylated and large parts of the methyltranscriptome are regulated by stress.** m^6^A-Seq of mouse cortex under basal conditions and 4 h after acute restraint stress (n = 7 each from mRNA of 3 mice pooled). **(B) Most m^6^A-peaks and stress-regulated m^6^A-peaks map to the coding sequence (CDS).** **(C) The peak distribution across the transcript length reveals enrichment** of m^6^A-peaks **at the 5’ UTR and the stop-codon.** Stress-regulated m^6^A-peaks show the same distribution as all m^6^A-peaks in both coding and non-coding transcripts. **(D) GGACWB is the most abundant motif detected in m^6^A-peaks and enriched at peak summits.** **(E) Stress-regulated peaks are equally distributed to up- and down-regulated peaks with the majority being altered in a quantitative fashion rather than being condition-specific.** **(F) Control-higher and control–specific peaks are enriched in 5’UTRs, whereas stress-higher and stress–specific genes are enriched in 3’UTRs.** See also Figure S1 G. **(G)Quantitative regulation of m^6^A by stress detected in m^6^A-Seq was replicated by m^6^A-RNA-immunoprecipitation (RIP)-qPCR in an unrelated cohort of animals.** Left panel per gene: averaged sequence tracks of 7 m^6^A-Seq biological replicates and peaks. Arrows indicate quantitatively regulated peaks with up/down arrows marking peaks significantly regulated with Q<0.05 and diagonal arrows marking peaks significantly regulated before multiple testing correction with P<0.05. Right panel per gene: differential peaks were validated in a separate cohort of mice using full length m^6^A-RIP-qPCR resulting in measurement of fold changes (FC) of m6A and mRNA. (n = 7, mean ± SEM, * depict omnibus Tukey post-hoc tests to basal P<0.05 after FDR-corrected one-way ANOVA. Con = Control, Str = Stress. Full statistics see Table S2. See also Figure S1 D) **(H) Stress-regulation of m^6^A negatively correlates with the respective change in mRNA.** (log2 fold changes plotted versus RNA fold changes stress to basal, m^6^A fold change is averaged in case of multiple peaks. n = 25,821, Generalised Linear Model see Table S2.) **(I) Genes with stress-related m^6^A-changes are enriched for genes bound by m^6^A-readers.** Mouse homologues of genes reported in previous PAR- and HITS-CLIP experiments in human cell lines were intersected with the genes harbouring differential m^6^A-sites. The amount of overlap observed (blue line) was compared to distributions gained from 100 random permutations (grey distributions) of all observed expressed genes with human homologues (Z-Test).

While the majority of peaks were located in the CDS (coding sequence) of transcripts (Figure 1 B), analysing the peak distribution along their length in respect to the landmarks of RNA transcripts (Cui et al., 2016) revealed an enrichment both at the 5’UTR (5’ untranslated region) and around the stop-codon (Figure 1 C). The m^6^A consensus motif DRACH and the more conservative motif GGAC were found in 93.36 % and 68.68 % of the m^6^A-peaks, respectively, often with several occurrences per peak sequence, indicating potential multiple adjacent m^6^A-sites. Sequence motif analysis confirmed enrichment of the m^6^A consensus motif, with the top motif being GGACWB (Figure 1 D). Functional classification of m^6^A-peaks harbouring transcripts revealed over-representation of genes related to neuronal plasticity, morphogenesis and development with different biological processes and pathways enriched depending on the peak’s position (Figure S1 C).

Differential analysis reported peaks that were either unique to the control- or the stress-condition as well as m^6^A-peaks with quantitative changes between the conditions (Figure 1 E). Interestingly, both control-unique and higher-in-control peaks exhibited position preference for the 5’UTR, whereas stress-unique and higher-in-stress peaks were enriched in the 3’UTR (Figure 1 F and Figure S1 E). Differential methylation of chosen candidate transcripts was confirmed by m^6^A-RNA immunoprecipitation (RIP) followed by qPCR (Figure 1 G and Figure S1 D). Notably, differential methylation of some genes that were removed from analysis because of their low expression could also be confirmed or at least showed a similar trend of regulation (Figure S1 C).

Although only a fraction of transcripts was significantly regulated both on the m^6^A and RNA-level after correction for multiple testing, changes in m^6^A across all genes negatively correlated with the respective change in transcript levels (Figure 1 H, Table S2). This may indicate that higher methylation levels may functionally lead to reduced RNA levels. The same negative relationship was detected for transcripts with stress-altered m^6^A (Table S2). Interestingly, the degree of the inverse relationship depended on the peak’s position, with CDS peaks showing the strongest negative correlations, followed by 5‘UTR peaks (Figure S1 F, Table S2).

Since the function of altered m^6^A likely involves the action of reader-proteins, we next asked if methylation-altered transcripts are preferentially recruited to m^6^A-binding proteins. When we intersected m^6^A-regulated genes with those bound by m^6^A readers, we observed a higher than expected overlap for several but not all m^6^A readers (Figure 1 I, Q (FDR-corrected P-Value) < 0.05 for YTHDF1, YTHDF2, YTHDC1, HNRNPC), indicating that stress-regulated m^6^A-peaks are poised for downstream functional targeting by m^6^A binding proteins. Additionally, analysing the co-occurrence of the observed m^6^A-motif GGACWB to known binding motifs of RNA-binding proteins within the m^6^A-peak sequences revealed a higher than expected overlap and summit enrichment with the binding motifs of FMR1, a protein crucial for directed translocation of RNAs within neurons and critical for synaptic plasticity (Figure S1 G). Likewise, genes reported to be bound by the mouse FMR1 or the human homologue FMRP were also higher-than-likely m^6^A-methylated (mouse FMR1 Z = 9.25, Q < 0.05; human FMRP Z = 62.9, Q < 0.05; Figure S1 H), suggesting that m^6^A methylation of neuronal RNAs may regulate protein binding critical for neuronal transport and plasticity.

### Stress-regulation of m6A is brain-region-specific

Based on the fact that the fold-changes of stress-regulated m^6^A-peaks were rather small in the mouse cortex m^6^A-Seq, we reasoned that the true extent of the m^6^A-stress response may only be revealed when investigating more defined brain regions. Therefore, we measured the time course of RNA methylation changes in 2 regions highly involved in stress response regulation: the medial prefrontal cortex (PFC) and the basolateral and central amygdala (AMY; Figure 2 A). We found that global m^6^A was regulated in total RNA in a region-dependent manner with RNA methylation decreased in the PFC and increased in the AMY (Figure 2 B). The same regulation was observed when m^6^A was measured in mRNA using LC-MS/MS (Figure 2 C). To examine potential changes of the m^6^A-machinery related to these global changes, we measured gene expression levels of m^6^A enzymes and binding proteins. We found the demethylases *Fto* and *Alkbh5* to be differentially regulated in a region-specific manner facilitating the effects observed on global methylation, in most cases preceding the effect observed on global m^6^A (Figure 2 D). Furthermore, *Mettl3* was downregulated upon stress-exposure tissue-independently (Figure 2 D) and *Wtap* was regulated isoform-specifically only in the AMY (Figure S2 B). The m^6^A-reader *Ythdc1* was regulated in a region-specific manner (Figure 2 D), whereas the other known enzymes and readers were not differentially expressed (Figure S2 A).

**Figure 2.**
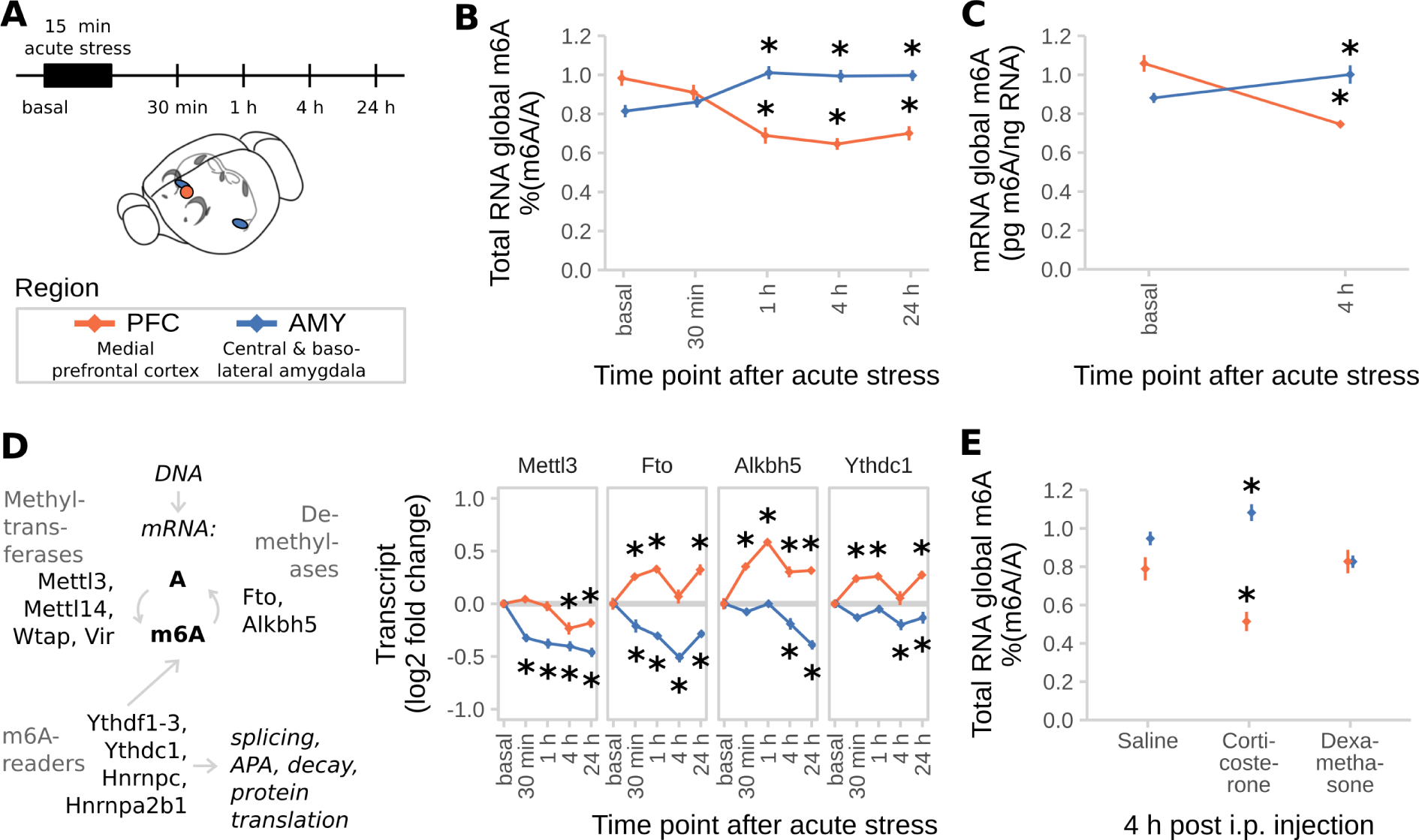
Acute restraint stress regulates brain global m6A and expression of the m6A regulatory system in a time- and region-specific manner. **(A) Experimental design. PFC = medial prefrontal cortex (orange), AMY = central and basolateral amygdala (blue).** **(B) Global m^6^A is decreased in the PFC and increased in the AMY after acute restraint stress.** (Global m^6^A assay on total RNA, n = 12, mean ± SEM. 2-way ANOVA interaction effect F(4,110) = 24.045, P<0.001. * depicts omnibus Tukey post-hoc tests to basal P<0.05. Results were replicated in 3 independent mouse cohorts with only 1 experiment shown). **(C) Likewise, global mRNA m^6^A is decreased when measured with LC-MS/MS.** (n = 7, mean ± SEM. 2-way ANOVA interaction effect F(1,24) = 159.537, P<0.001. * depicts omnibus Tukey post-hoc tests to basal P<0.05). **(D) m^6^A regulatory genes *Mettl3*, *Fto*, *Alkbh5* and *Ythdc1* are differentially expressed after acute stress in the brain.** For further m^6^A-related genes see Figure S2. (n = 12, log2 fold change ± SEM. 2-way MANOVA: significant interaction effects for *Fto*, *Alkbh5*, and *Ythdc1*, main stress effect for *Mettl3*, each FDR-corrected P< 0.05 and n_2_>0.01. * depicts omnibus Tukey post-hoc tests to basal P<0.05. Full statistics see Table S2. Results were replicated in 3 independent mouse cohorts with only 1 experiment shown). **(E) Global m^6^A is decreased in the PFC and increased in the AMY after corticosterone i.p. injection, but not after dexamethasone injection.** Corticosterone: 250 µg/kg, dexamethasone 10 mg/kg. (Global m^6^A assay on total RNA, n = 12, mean ± SEM. 2-way ANOVA reported a significant interaction effect (F(4,96) = 12.887, P<0.001). * indicates omnibus Tukey post-hoc tests P<0.05 compared to area basal).

Notably, i.p. injection of corticosterone but not dexamethasone changed global m^6^A (Figure 2 E) as well as *Fto* and *Alkbh5* expression (Figure S2 B) similarly to acute stress (Figure 2 D), demonstrating that the stress effect may be mediated by glucocorticoids (GCs). Supporting this idea, we found that the majority of m^6^A enzyme- and reader-genes contains several GC response elements in their 5’ upstream region, likewise pointing at expression regulation of those genes via GCs (Figure S2 C).

### Stress-regulation of m6A is gene-specific

m^6^A-Seq requires large amounts of input material thus limiting the capacity of the technique to investigate narrowly defined brain regions and also only measures relative enrichment changes and not absolute transcript methylation. Therefore, we performed m^6^A-immunoprecipitation of full length transcripts (m^6^A-RIP) followed by qPCR to assess absolute levels of candidate transcript methylation in narrowly defined brain areas, before and after stressful challenge. For calibration of the assay and normalization of immunoprecipitation-efficiency in downstream experiments, we designed and used an m^6^A-methylated internal spike-in RNA oligonucleotide (Figure 3 A and Figure S3 A, B). The m^6^A-RIP-qPCR detected m^6^A methylated RNA spike-in across a wide range of concentrations with low IgG-background signal and without competing with the immunoprecipitation of endogenously methylated RNAs (Figure 3 B). Using mixtures of unmethylated and methylated spike-in oligonucleotides, we confirmed that m^6^A-RIP-qPCR measured different methylation states of RNAs with high precision (Figure 3 C, r²>0.95).

**Figure 3.**
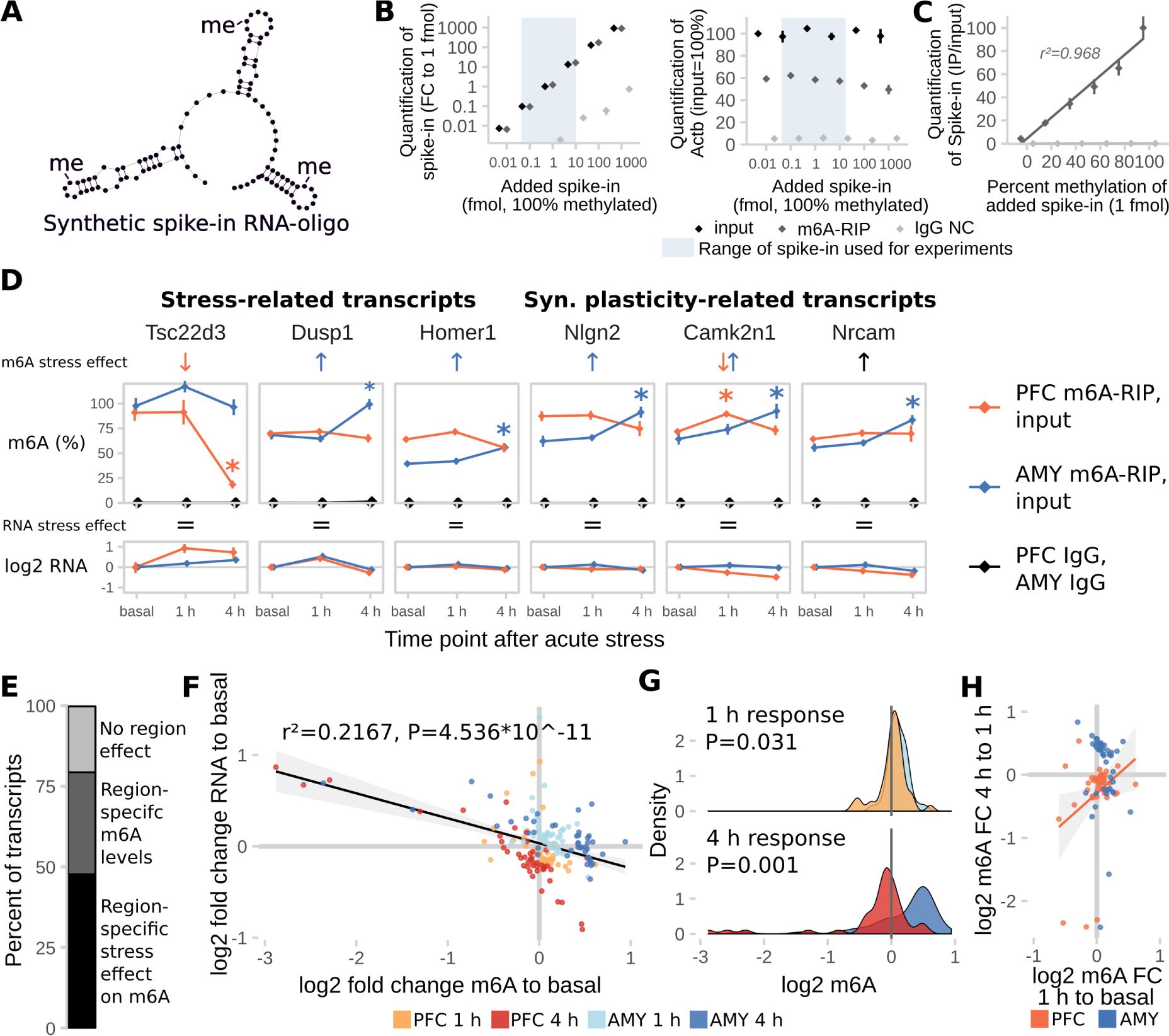
Absolute regulation of m6A methylation is site-specific. **(A) A synthetic RNA oligonucleotide with 3 internal m^6^A-sites was used for validation and internal normalization of the m^6^A-RIP-qPCR.** For additional characterization of the RNA oligonucleotide see also Figure S3. **(B) m^6^A-RIP-qPCR detects the methylated spike-in oligonucleotide in a linear fashion without impairing precipitation-efficiency for endogenous transcripts in the concentration range used for experiments.** Fully methylated spike-in oligo was added to unfragmented total RNA and precipitated with anti-m^6^A antibody (m^6^A-RIP) or rabbit IgG (IgG NC) (n = 3 technical replicates, normalized expression to 1 fmol input control. Mean ± SEM). **(C) m^6^A-RIP-qPCR accurately quantifies differential methylation of the spike-in oligo.** 1 fmol spike-in oligo mixed from fully methylated and fully unmethylated spike-in was added to unfragmented total RNA and precipitated with m^6^A-RIP-qPCR (n = 3 technical replicates, normalized to input control. Mean ± SEM). **(D) Absolute full length m^6^A-levels of stress-related and synaptic plasticity-related transcripts are differentially regulated in the PFC and AMY of stress-related candidate transcripts and synaptic-plasticity-related candidate transcripts after stress.** See also Figure S3. (n = 8, mean ± SEM. Significant effects observed in FDR-corrected 2-way MANOVA (P<0.05, n²>0.01) are coded in the rows “m^6^A stress effect” and “RNA stress effect”: orange/blue arrows = PFC-/AMY-specific stress effect (interaction effect 2-way ANOVA, one-way follow up significant in respective tissue), black arrow = stress main effect, equals sign = no interaction or stress main effect in 2-way ANOVA. See also Table S2). **(E) The majority of transcripts measured are expressed or regulated in a region-specific manner.** (Percent of transcripts with significant interaction or main effect in FDR-corrected 2x2 MANOVA). **(F) Stress-regulation of m^6^A negatively correlates with changes in RNA levels.** (log2 fold changes of m^6^A and RNA after stress to basal time points, n = 44 per group, black line: Linear model + 95% CI. GLMs see Table S2). **(G) General patterns of m^6^A-changes vary in extent and direction depending on brain region and time point.** (Density plots of data depicted in (D), T-Test). **(H) The m^6^A change at the 1 h time point correlates with the m^6^A change at 4 h in the PFC, but not AMY, indicating that in the PFC, m^6^A change 1 h after stress is a proxy for later change.** (Orange line: Linear model for PFC only + 95% CI. GLMs see Table S2).

Applying m^6^A-RIP-qPCR, we measured absolute methylation levels of several candidate transcripts involved in the brain’s stress response and, given the enrichment of neuronal plasticity and morphogenesis-related terms in the m^6^A-Seq, synaptic plasticity-related transcripts (Figure 3 D and Figure S3 C). Similar to the results of the m^6^A-Seq, regulation of m^6^A by stress (26/44 transcripts) was observed more often than regulation of RNA (16/44 transcripts, with 12 overlapping) in the transcripts tested. Notably, the majority of chosen candidates were either regulated or expressed in a region-specific manner, emphasizing the importance of assessing RNA-methylation in defined brain areas (Figure 3 E). The negative correlation of m^6^A- and RNA fold-changes was replicated in the m^6^A-RIP-qPCR data, with no influence of region and time point (Figure 3 F, Table S2). In detail, both PFC and AMY exhibited differential response both at 1 h and 4 h with opposite directions, paralleling the regulation observed in global m^6^A in the respective regions before (Figure 2 A). Overall, 4 h fold changes had higher effect sizes compared to 1 h fold changes (Figure 3 G). Fold changes at the 1 h time point correlated with those at 4 h for the same gene in the PFC but not AMY indicating that in the PFC 1 h m^6^A may be an intermediate state of 4 h regulation with fold-changes of regulated m^6^A increasing with time. In contrast, in the AMY for the candidate genes investigated, m^6^A regulation after 1 h and 4 h was more independent (Figure 3 H).

### Stress-coping behaviour is altered in mice deficient of *Mettl3* or *Fto*

To explore the causal consequences of disturbed m^6^A levels on the stress response and stress-linked behaviours, we investigated mouse models of altered m^6^A-regulation in adult neurons. Since the expression of the m^6^A methyltransferase *Mettl3* and the m^6^A and m^6^Am demethylase *Fto* were affected by acute stress, we generated conditional inducible knockout mouse models lacking these genes specifically in forebrain excitatory neurons by breeding *Mettl3* or *Fto* flox/flox mice to tamoxifen-inducible Nex-CreERT2 mice (Mettl3-cKO and Fto-cKO). Upon induction with tamoxifen in young adults to prevent developmental effects, *Mettl3* and *Fto* were depleted both from dorsal and ventral parts of the hippocampus, specifically in CA1 and CA3 but not in the dentate gyrus (Figure 4 A). Nex-CreERT2-induced recombination is further known to occur in small populations of principal neurons in the

**Figure 4.**
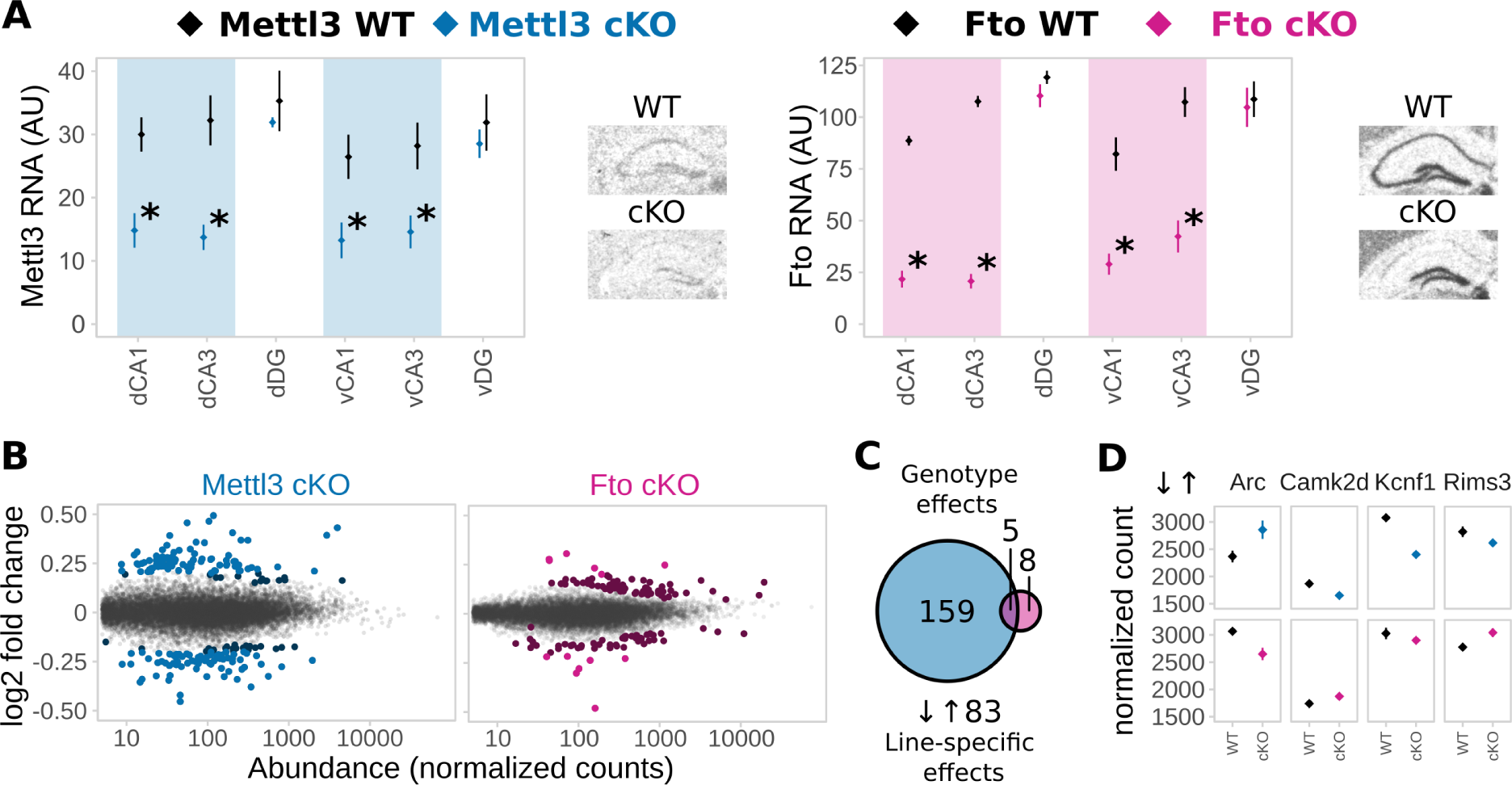
Deletion of *Mettl3* or *Fto* in adult excitatory neurons of the hippocampus CA1 and CA3 alters gene expression in animals. **(A) *Mettl3* and *Fto* are depleted from the dorsal (D) and ventral (v) hippocampus CA1 and CA3 in Mettl3-cKO (blue) and Fto-cKO (pink) mice, respectively.** WT = wild type, cKO = conditional knockout, DG = dentate gyrus. RNA expression of the loxP-targeted gene exon was measured by *in-situ*-hybridization. (Optical density normalized to background in arbitrary units (AU); mean ± SEM. n = 4, signal averaged across both hemispheres, * depict T-Tests P<0.05). **(B) mRNA-Seq of adult CA1 and CA3 shows altered gene expression after deletion of *Mettl3* and *Fto* in non-stressed basal animals.** (Scatter plots of log2 change ratio to DeSeq normalized count gene abundance of all transcripts detected in the RNA-Seq of adult basal animals. Differentially expressed genes are indicated in blue (Mettl3-cKO) and pink (Fto-cKO) with Q<0.1, with darker colours marking genes that were differentially expressed but had fold changes below the set cut-off of absolute log2 fold change>0.2. n=5) **(C) More genes are differentially expressed after deletion of *Mettl3* (164) compared to deletion of *Fto* (13) with very few overlapping (5).** 83 genes are expressed in a knockout x genotype-specific pattern (Q<0.1, absolute log 2 fold change >0.2) (D) 4 representative examples of genes expressed in a knockout X genotype specific pattern. (Genotype effects shown for *Mettl3* (blue) or *Fto* deletion (pink). Line-specific effects depict knockout x genotype interaction effects. Normalized counts indicate DeSeq2 normalized counts. n=5)

cortex (*33*). Depletion of either gene did not result in compensatory changes of gene expression of other genes involved in m6A-metabolism (Suppl. Figure S4 A), but altered transcriptome profiles as observed by mRNA-Seq of CA1 and CA3 tissue (Fig 4 B). Interestingly, in non-stressed basal animals, we observed a larger number of differentially expressed genes in Mettl3-cKOs compared to Fto-cKOs (Figure 4 B, C; Mettl3-cKOs: 205 differentially expressed genes with 164 genes with a fold change above log2=0.2; Fto-cKOs: 130 differentially expressed genes with 13 genes with a fold change above log2=0.2; Suppl. Table 3) as well as a wider spread of gene expression regulation (Supp. Figure S4 B) with no apparent preference for up- or downregulation. Although there was only small overlap of differential expressed genes between the two lines, 91 genes were differentially expressed in a knockout-specific pattern (Figure 4 C; 83 genes with a significant interaction effect of gene knockout and genotype and a fold change above log2=0.2; Suppl. Table 3) including crucial genes for regulation of neuronal activity response, and synaptic function as immediate-early genes as well as genes involved in synaptic functions (Figure 4 C). Neither Mettl3-cKO nor Fto-cKO mice showed altered anxiety-like behaviour or locomotion (assessed by the Open Field, Elevated Plus Maze and Dark-Light-Box tests; Figure S5 A) but we observed significant changes in spontaneous digging behaviour (Marble Burying Test, Figure S5 A). Both mouse models exhibited increased cued fear memory long-term maintained during memory extinction (Figure 5 A), suggesting that both perturbations in the m^6^A-system lead to long-term increased fear memory. Additionally, Fto-cKO mice exhibited increased contextual fear memory (Figure 5 A). Importantly, we observed no differences in non-fear-related memory or short-term working memory when tested in the object recognition test and Y-Maze spontaneous alternation test (Figure 5 A). Next, we investigated the transcriptional response patterns 24 h after fear conditioning stress, thus at the time point we observed the altered memory, comparing fear conditioned animals (“FC”) to control animals that experienced the same handling but no foot shock (“Box”). For both Mettl3-cKOs and Fto-cKOs we observed a large number of genes differentially expressed after fear conditioning in a genotype-dependent manner, implying a widely altered transcriptional response pattern after stress in animals with disturbed m6A-system (Figure 5 B). Thereby, significant gene-expression regulation was more extended in fear conditioned animals compared to non-fear conditioned animals (supp. Figure S4 C; Suppl. Table 3) with low overlap between both mouse models (Figure 5 C). In contrast to basal animals, Fto-cKOs showed more genotype-dependent expression changes after the stressful fear conditioning event than Mettl3-cKOs (Figure 5 B; Suppl. Table 3), implying that despite the major lack of Fto-cKO genotype effect in steady-animals, *Fto* is crucial for the regulation of the stress and fear response. Upon fear-conditioning stress, genotype-dependent transcriptomic changes involve genes crucial for neuronal systems like neurotransmitter receptors and transporters as well as transcription factors (Figure 5 C), pointing at a role of m^6^A in regulating synaptic function in the hippocampus after fear conditioning. Consequently, investigating the effects of *Fto* and *Mettl3* depletion on electrophysiological correlates of network plasticity and brain function, we found that CA1 long-term potentiation was impaired in Fto-cKO but not in Mettl3-cKO mice (Figure 5 D) with no effect on paired-pulse facilitation (Figure 5 D) nor basal neurotransmission (Figure S5 B).

**Figure 5.**
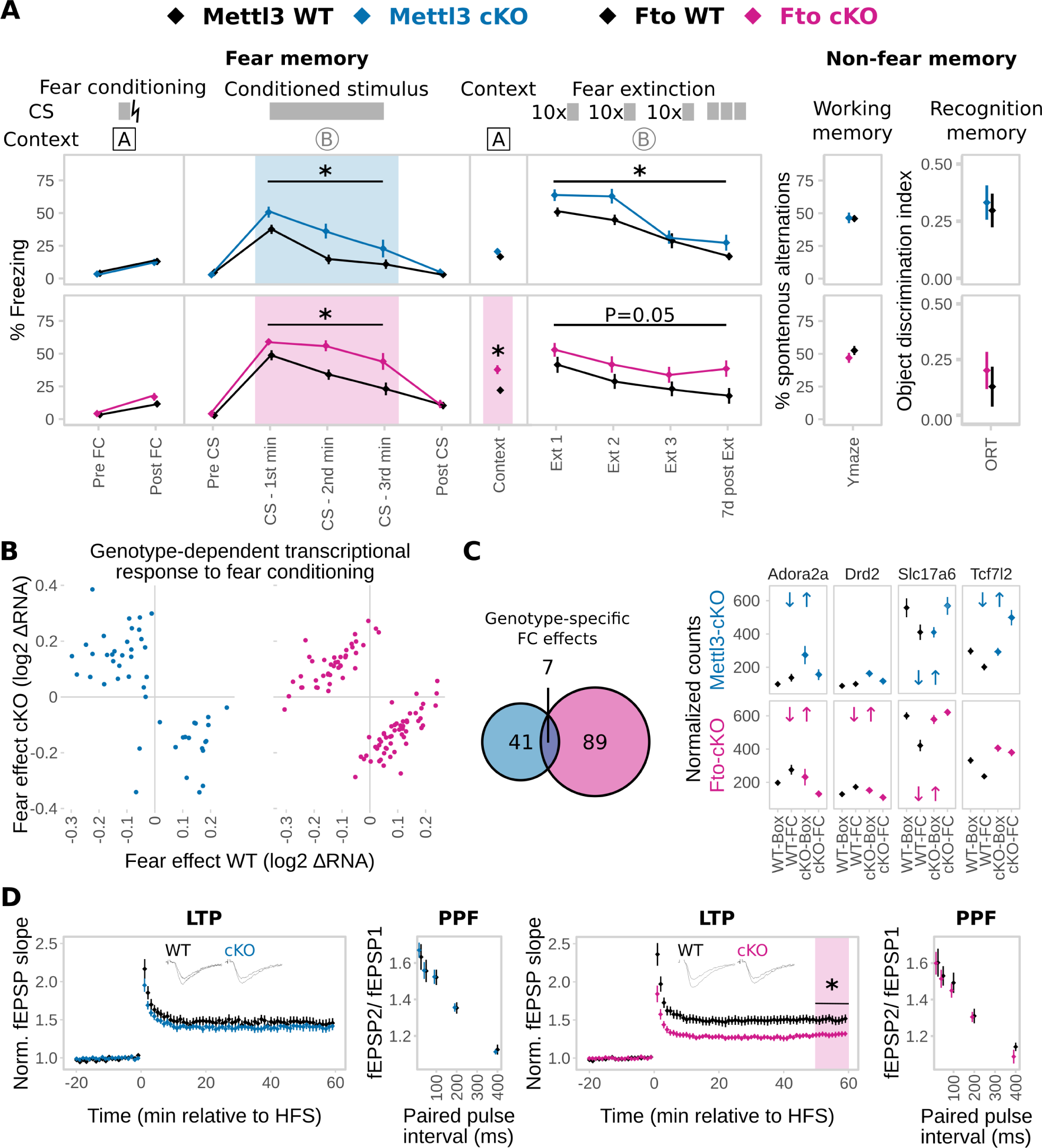
Animals with adult-excitatory-neuron-specific depletion of *Mettl3* and *Fto* showed impaired fear coping potentially mediated by a differential transcriptomic response to fear and encoded by changes in hippocampus CA1 electrophysiological properties. **(A) Both Mettl3-cKO (blue) and Fto-cKO (pink) animals display increased conditioned fear memory long-term maintained during fear extinction.** The primary fear response was not altered. Fto-cKO animals also have increased contextual fear memory. No difference was observed in the Y-Maze test or the Object-Recognition-Test (ORT). CS = Conditioned Stimulus, lightning bolt = US = unconditioned stimulus. Ext = Extinction. (n = 11–13, mean ± SEM. Fear expression was binned in 1 min intervals during CS representation. * depicts a main genotype effect in repeated measurements ANOVA for CS and Ext bins and a T-Tests P<0.05 for all other data points). **(B) The transcriptomic response 24 h after fear conditioning (FC) is altered in both animals with *Mettl3* (blue) or *Fto* (pink) depletion.** (Scatter plot of log2 RNA fold change in WT vs. cKO animals of only those genes with a significant genotype x FC effect. Q<0.1, absolute log 2 fold change >0.2, n=5) **(C) More genes express a genotype-dependent FC-effect in Fto-cKOs compared to Mettl3-cKOs with low overlap.** 4 examples of such genes are shown. (Significant genotype x FC in the examples are depicted by blue (Mettl3-cKOs) and pink (Fto-cKOs) opposite arrows. Q<0.1, absolute log 2 fold change >0.2, n=5) **(D) Long-term potentiation (LTP) but not short-term plasticity in CA1 was attenuated in Fto-cKO mice (pink) but not Mettl3-cKO mice (blue).** Short-term synaptic plasticity was measured by paired-pulse facilitation (PPF). (n = 10–12 slices from 5–6 animals, mean ± SEM plus representative LTP trace curves, HFS = high frequency stimulation. * Depicts T-Test P<0.05 on the average field excitatory postsynaptic potential (fEPSP) slope 50–60 min post HFS).

### Regulation of m6A is impaired in MDD patient blood

To evaluate the potential of blood m^6^A as a peripheral proxy of the central m^6^A stress response, we measured global m^6^A methylation levels in mouse and human blood after an acute stressful challenge and GC stimulation. Global methylation was transiently decreased in whole blood of mice after acute stress (Figure 6 A), with gene expression of *Mettl3* and *Alkbh5* altered in accordance with the global m^6^A change and *Wtap* being upregulated (Figure 6 B). Similarly, global m^6^A was decreased in mouse blood 4 h after i.p. injections of both corticosterone and dexamethasone (Figure 6 C). Comparably, blood from healthy human volunteers, drawn before and after intake of 1.5 mg dexamethasone, showed both reduced global m^6^A levels (Figure 6 D) and changes in gene expression of the m^6^A-machinery enzymes 3 h after dexamethasone intake (Figure 6 E) (Arloth et al., 2015).

**Figure 6.**
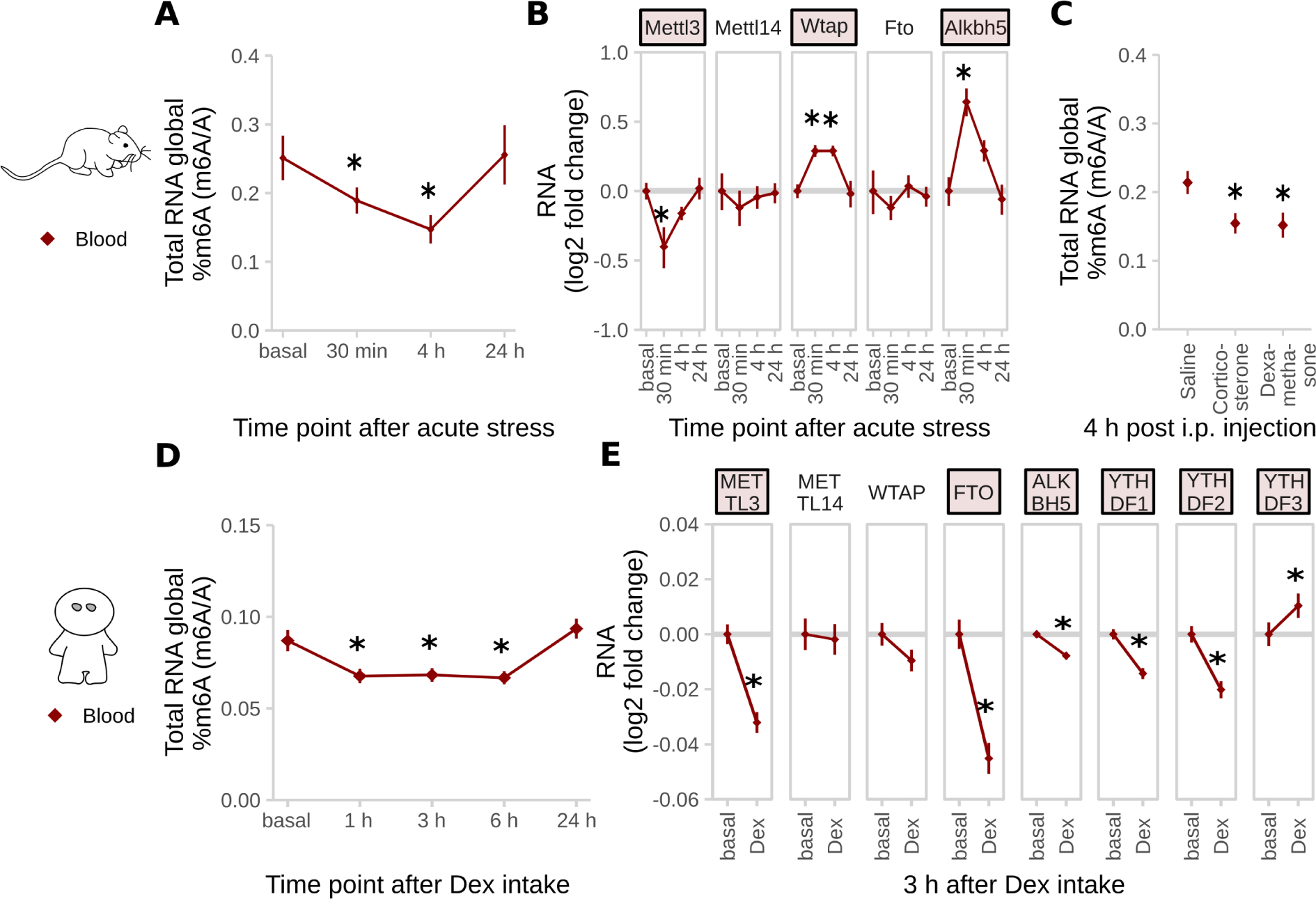
Global m6A in the blood is transiently decreased after stress in mice and stimulation with glucocorticoids (GCs) in humans. **(A) Global m^6^A is transiently decreased in mouse blood after acute stress.** (Global m^6^A assay on total RNA, n = 8, mean ± SEM. * depict omnibus post-hoc comparisons to basal, P<0.05, after Kruskal-Wallis-Test P<0.05). **(B) Global m^6^A changes in mouse blood are accompanied by changes in m^6^A regulatory genes.** (qPCR on total mouse blood, log2-fold changes of different genes to basal. n = 8, mean ± SEM, Red coloured gene names: one-way ANOVA, * depict omnibus Tukey post-hoc tests to basal P<0.05, see Table S2). **(C) Furthermore, global m^6^A is decreased in mouse blood both after corticosterone and dexamethasone i.p. injection.** Corticosterone: 250 µg/kg, dexamethasone 10 mg/kg. (Global m^6^A assay on total RNA, n = 12, mean ± SEM. 2-way ANOVA reported a significant interaction effect (F(4,96) = 12.887, P<0.001). Stars indicate omnibus Tukey post-hoc tests P<0.05 compared to area basal). **(D) In a similar way, global m^6^A is temporarily decreased in the blood of healthy human subjects after treatment with 1.5 mg dexamethasone (Dex).** (Global m^6^A assay on total whole blood RNA, n = 25 healthy men, mean ± SEM. Kruskal-Wallis-Test P<0.001, * depict omnibus Tukey post-hoc tests to basal P<0.05) **(E) Expression of m^6^A regulatory genes in human blood is also affected by dexamethasone.** (Human whole blood at baseline and 3 H after intake of blood, data extracted from microarray (Arloth et al., 2015), n = 160 healthy and diseased subjects, mean ± SEM, * depict Bonferroni-corrected T-Tests to basal P<0.05). See also Figure S4.

Since dysregulation of the stress response may be an important feature of psychopathologies like Major Depressive Disorder (MDD), we next investigated whether m^6^A-regulation in response to dexamethasone differs between healthy individuals and MDD patients. In contrast to healthy subjects, down-regulation of m^6^A in response to dexamethasone was neither observed in male nor female MDD patients (Figure 7 A). To prevent potential contamination of results by antidepressant treatment present in blood of MDD patients, we confirmed the lack of response to GC-stimulation using dexamethasone (Figure S5 A) and cortisol (Figure 7 B total RNA, Figure 7 C mRNA by LC/MS-MS) in B-lymphocyte cell lines (BLCLs) donated by each 6 healthy volunteers and 6 MDD-patients propagated in absence of antidepressants. NR3C1 (GC Receptor) mRNA and protein expression as well as transcriptional response to GC stimulation was unchanged in BLCLs of MDD donors (Figure S6 B-d). To estimate the transcriptome-wide distribution of m^6^A after GC stimulation, we performed m^6^A-Seq on BLCLs of one healthy donor treated for 1 h with 100 nM cortisol or mock conditions. We found over 17,000 m^6^A-peaks in around 9,000 genes in the healthy donor BLCLs (89% of peaks with GGAC motif, 99% of peaks with DRACH motif), with 12 % of those being cortisol-treatment-responsive (Figure 7 D, regulated peaks with absolute log2 fold change > 1, FDR-corrected P-Value < 0.05). Similar to the mouse m^6^A-Seq, we found the m^6^A consensus motif was enriched in the m^6^A-peaks (top motif GKACW) and m^6^A-peaks were distributed along the transcript-length with preference for both m^6^A-peaks in the 5’UTR and around the stop codon. Interestingly, in m^6^A-peaks that were found to be cortisol-responsive, we observed less preference for the 5’UTR with higher contribution of CDS and 3’UTR peaks (Figure 7 E,F). In parallel to the global demethylation observed in BLCLs of healthy donors after cortisol-stimulation, differential m^6^A-peaks were found to be almost exclusively hypomethylated with higher fold changes than observed in mouse brain (Figure 7g). To precisely quantify regulation of m^6^A-levels in BLCLs from healthy and MDD-donors, we performed m^6^A-RIP-qPCR testing for GC-responsive genes in BLCLs after stimulation with cortisol. We observed specific downregulation of m^6^A in *FKBP5*, *IRS2* and *TSC22D3* in cells from healthy but not from MDD individuals (Figure 7h). In line with the general trends observed before, methylation of tested candidates in cells derived from healthy, but not MDD donors, was significantly decreased (Figure 7i).

**Figure 7.**
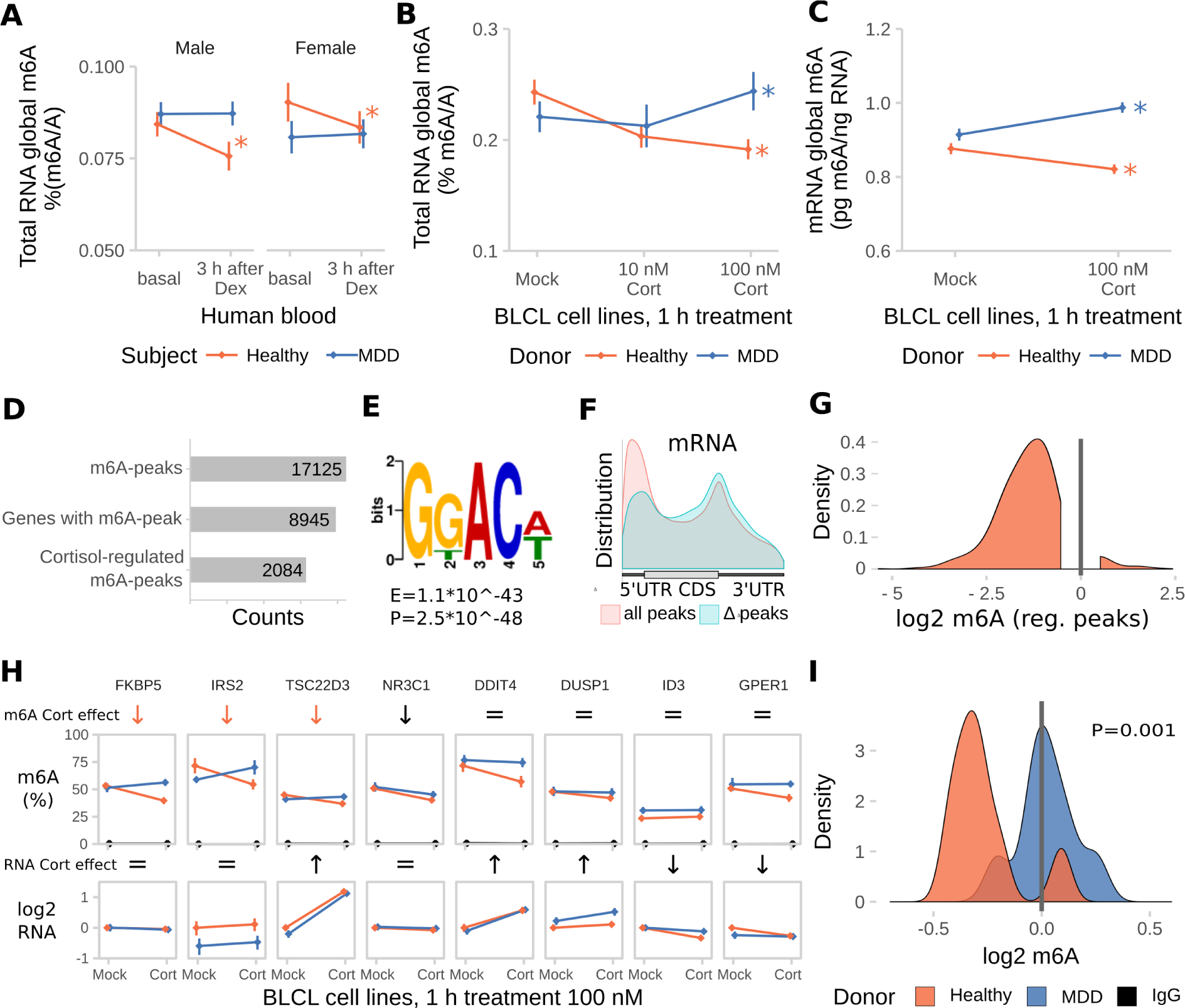
The glucocorticoid induced m6A-reduction in blood is absent in blood and cell lines from donors with Major Depressive Disorder (MDD). Dex = Dexamethasone, Cort =Cortisol. **(A) The dexamethasone induced m^6^A decrease in human blood m^6^A is absent in MDD patients.** (Male and female, healthy and MDD subjects each, n = 25, mean ± SEM. 3-way mixed-design ANOVA: significant interaction effect of treatment and subject status (F(1,96) =11.184, P = 0.001), but no interaction with sex. * depict omnibus Tukey post-hoc tests to sex basal P<0.05). **(B) Global m^6^A is decreased in B-lymphocyte cell lines (BLCLs) in a concentration dependent manner after 1 h treatment with cortisol**. (Global m^6^A assay on total RNA, n = 5 biological replicates with 3 technical replicates each, mean ± SEM. 2-way ANOVA: significant interaction effect of cortisol and donor status (F(3,24) = 44.365, P<0.001). * depict omnibus Tukey post-hoc tests to basal P<0.05). **(C) The same regulation is observed on mRNA using LC-MS/MS.** (n = 5, mean ± SEM. 2-way ANOVA significant interaction effect of cortisol and donor status (F(1,20) = 19.196, P<0.001). * depict omnibus Tukey post-hoc tests to mock treatment P<0.05). **(D) m^6^A-Seq reveals large proportions of the BLCL transcriptome to be methylated and potentially stress-responsive.** (m^6^A-Seq of one BLCL cell line of a healthy donor treated for 1 h with 100 nM cortisol or mock.) **(E) Similar to the mouse brain, the m^6^A consensus motif is enriched in BLCL m^6^A-peaks (top motif GKACW).** **(F) BLCL-m^6^A-peaks show a similar distribution across the transcript length like the mouse brain with less contribution of 5’UTR peaks for cortisol-responsive genes.** **(G) In accordance with global demethylation after cortisol-stimulation in BLCLs of healthy donors, the majority cortisol-responsive m^6^A-peaks were found to be downregulated by cortisol in m^6^A-Seq.** (Distribution of log2 fold change of m^6^A-enrichment signal of identified cortisol-responsive m^6^A-peaks). **(H) Cortisol-responsive genes FKBP5, IRS2 and TSC22D3 m^6^A are specifically downregulated in cell lines of healthy, but not MDD donors after stimulation with cortisol, when assessed with m^6^A-RIP-qPCR.** (m^6^A-RIP-qPCR. n = 5, mean ± SEM. Significant effects observed in FDR-corrected 2-way MANOVA (P<0.05) are coded in the rows “m^6^A Cort effect” and “RNA Cort effect”: orange arrows = healthy donor-specific Cort effect (interaction effect 2-way ANOVA, one-way follow up significant in healthy donors only black arrow = Cort main stress effect, equals sign = no interaction or stress main effect in 2-way ANOVA. Full statistics see Table S2). **(I) Density plots m^6^A change upon cortisol treatment.** (Density plots of log2 fold change data as m6A-RIP-qPCR-data depicted in (G), donor-dependent-distributions of fold changes were compared using a T-Test).

## Discussion

Here, we have identified m^6^A and m^6^Am as a widely regulated epitranscriptomic mark responsive to acute stress. Using m^6^A-Seq in mouse cortex, we observed over 25,000 m^6^A-peaks with around 7% being stress-responsive, thus far outnumbering the changes observed on the RNA expression level. Peaks were enriched at both the 5’UTR and around the stop-codon with a higher peak contribution in the 5’UTR than previously reported (Meyer et al., 2015). In contrast to previous reports indicating changes in peaks primarily at the 5’UTR in cellular heat shock response (Zhou et al., 2015), we observed no preference in location for differential peaks. However, we detected a preference for stress-downregulated peaks for the 5’UTR and for stress-upregulated peaks for the 3’UTR. Given the wide-variety of mechanisms of how RNA methylation may regulate RNA levels and translation, the molecular consequence of differential m^6^A regulation after stress is not yet clear. Stress-m^6^A regulated genes were significantly enriched for target genes of several m^6^A-readers as well as for the neuronal RNA-binding and cell-transport-regulating protein FMR1/FMRP recently shown to bind m^6^A (Edupuganti et al., 2017), suggesting that the effects of altered methylation may be equally broad. Overlap of FMRP-binding sites with synaptic transcripts bound by NSun2, a methyltransferase for another RNA modification, has been suggested before (Hussain and Bashir, 2015). Although we observed a general negative correlation between m^6^A-change and RNA-abundance, most m^6^A-changes were not accompanied by significant transcript changes neither in m^6^A-RIP-qPCR nor in m^6^A-Seq. This may imply that differential m^6^A may act both by regulating RNA decay as well as location and translation control. Our m^6^A-Seq reported relatively small fold changes likely due to the large cellular heterogeneity of the material used with only a small fraction of cells being responsive to the treatment. Further, we propose that m^6^A regulation is highly specific to smaller brain areas with even potential opposite regulation in different areas leading to an underestimation of the actual regulation in m^6^A-Seq. This was supported by region-specific investigation of global and gene-specific m^6^A levels in the PFC and AMY. These 2 areas regulate behavioural and hormonal stress responses, fear and anxiety (McEwen et al., 2015), with the PFC exhibiting top-down control of the AMY in anxiety and fear in mice (Adhikari et al., 2015). These changes were accompanied by matching regulation in the demethylase *Fto* and *Alkbh5* expression, as well as regulation of the methyltransferase *Mettl3*. Interestingly, previous reports also showed transcriptional regulation of *Fto* after acute stress by fear conditioning (Walters et al., 2017; Widagdo et al., 2016). Further, we observed regulation of the direct m^6^A-reader Ythdc1 after stress. In Drosophila, the *Ythdc1* homologue *YT521-B* regulates neuronal function with behavioural defects in knockout-flies (Lence et al., 2016). Notably, we observe that corticosterone i.p. injection in mice causes similar effects on m^6^A and enzyme-expression like acute stress, pointing towards a potential signalling mechanism via centrally acting GCs. Additional work is needed to unravel the pertinent signalling cascades involved. Single-cell RNA-Seq data showed that all m^6^A enzymes and readers are expressed in all major brain cell types including glia (Zeisel et al., 2015). Therefore, it is unclear which of these cell types drive the observed expression changes. To investigate the mechanisms of m^6^A-methylation in neurons only, we specifically depleted *Mettl3* and *Fto* from adult excitatory neurons of the hippocampus and cortex. We observed increased fear memory for both cued and contextual fear (latter in Fto-cKO mice only), which may be indicative of impaired stress-coping. Increased fear expression was previously reported upon knock-down of *Fto* in the dorsal hippocampus (Walters et al., 2017) and in the PFC (Widagdo et al., 2016). Extending the previous findings, we found fear memory to be enhanced upon disruption of both m^6^A methyltransferase *Mettl3* or demethylase *Fto*, indicating that fear-memory requires fine-tuned regulation of m^6^A-levels rather than being directly regulated by m^6^A levels. Anxiety-like behaviour was not altered in either mouse line but may be regulated by m^6^A affected by different enzymes or in different brain areas and cell types other than those investigated here. Importantly, we did not observe other impairments in cognition or memory in these mice, pointing towards a potential importance of m^6^A for fear-specific memory. Mechanistically, we found that both *Mettl3* and *Fto* depletion not only alter the steady-state transcriptome in adult hippocampal neurons but also the transcriptomic response to the fear conditioning stress, including regulation of several genes involved in neuronal circuits function pointing out a function of m^6^A in regulating neuronal circuits. Consequently, we describe that network plasticity is specifically altered in the CA1, a brain region crucial for contextual fear, in Fto-cKO, but not Mettl3-cKO mice. This may reflect a neuronal correlate of altered m^6^A underlying the altered contextual fear memory observed in Fto-cKO mice. Several arguments point at both enzymes targeting largely non-shared methylation sites and/or genes in the specific case investigated, either by different m^6^A sites affected or by *Mettl3* targeting predominantly m^6^A and *Fto* predominately targeting m^6^Am: First, we did not observe oppositely directed behavioural or electrophysicological effects in both lines. Second, the gene sets affected in transcription by *Mettl3* and *Fto* depletion were widely non-overlapping. Lastly, both enzymes showed different contribution to steady-state transcriptomic regulation (mainly Mettl3-cKOs) and fear conditioning-regulated gene expression (dominantly Fto-cKOs).

Finally, we propose that regulation of m^6^A and its cellular machinery in blood may represent a peripheral proxy for part of the brain’s m^6^A response, similar to DNA methylation changes (Ewald et al., 2014; Provençal et al., 2012). Both mice and humans showed global blood demethylation after stress or GC intake, respectively. The m^6^A-response to GCs is impaired in blood and blood cells obtained from MDD patients, which may be a consequence of the altered GC receptor reactivity and downstream signalling reported in MDD (de Kloet et al., 2005). Interestingly, genetic variants in *FTO* (Milaneschi et al., 2014; Samaan et al., 2013) and *ALKBH5* (Du et al., 2015) have been reported to associate with risk for MDD before but are yet to be replicated in larger cohorts. There is growing evidence that fine-tuned transcriptional regulation is especially relevant for psychiatric disorders including disease-associated SNPs in enhancer regions (Cross-Disorder Group of the Psychiatric Genomics Consortium, 2013) and epigenetic changes (Klengel and Binder, 2015) including regulation of chromatin conformation (Won et al., 2016), histone modifications (The Network and Pathway Analysis Subgroup of the Psychiatric Genomics Consortium, 2015) as well as short and long ncRNAs (Issler and Chen, 2015; Parikshak et al., 2016). Here, we reveal RNA modifications as a novel layer of regulation of gene expression that is potentially crucial for these disorders. Integrating all layers of transcriptional regulation will be highly relevant for understanding psychiatric disorders.

In summary, m^6^A and m^6^Am methylation constitute a novel layer of complexity in gene expression regulation following stress exposure, which is pivotal for the adaptation of stress-responsive circuits to acute challenges. The exciting finding of m^6^A dysregulation in MDD opens the possibility for the development of novel diagnostic biomarkers and eventually to better treatments for anxiety disorders, depression and other stress-related diseases.

## Author contributions

M. En., and A.C. conceived and designed the experiments and wrote the manuscript. M.En. performed, and analysed most experiments. M. En. and S.R. performed bioinformatics analysis. C. E., P. M. K, L.T., and M.R.-H. assisted in experiments. M. J. performed preliminary experiments with no data used. M. U. performed and analysed mass spectrometry experiments. M. Ed. performed and analysed electrophysiological studies. J.A. and E. B. B. provided human microarray data. P.W., C. T. W., M. V. S., J. M. D., E. B. B., and A.C. conceived and designed the project. The project was supervised by A.C.

## Acknowledgements

We thank Christian Namendorf, Tamara Gerlach, Carine Dournes, Eva-Maria Wagner, Barbara Hauger, Ania Mederer, Andrea Ressle, Andrea Parl, and Daniela Harbich for their technical support and Albin Varga and the animal care team for their devoted assistance with animal care. We thank Stoyo Karamihalev, Michaela Filiou, Andreas Menke, and Andreas Genewsky for helpful discussions and Jessica Keverne for professional English editing. We thank Gidi Rechavi, Sharon Moshitch-Moshkovitz and Vera Hershkovitz for help with preparing m^6^A-Seq libraries and helpful discussions.

A.C. is the head of the Max Planck Society - Weizmann Institute of Science Laboratory for Experimental Neuropsychiatry and Behavioral Neurogenetics. This work was supported by: an FP7 Grant from the European Research Council (260463, A.C.); a research grant from the Israel Science Foundation (1565/15, A.C.); the ERANET Program, supported by the Chief Scientist Office of the Israeli Ministry of Health (A.C.); the project was funded by the Federal Ministry of Education and Research under the funding code 01KU1501 A (A.C.); research support from Roberto and Renata Ruhman (A.C.); research support from Bruno and Simone Licht; I-CORE Program of the Planning and Budgeting Committee and The Israel Science Foundation (grant no. 1916/12 to A.C.); the Nella and Leon Benoziyo Center for Neurological Diseases (A.C.); the Henry Chanoch Krenter Institute for Biomedical Imaging and Genomics (A.C.); the Perlman Family Foundation, founded by Louis L. and Anita M. Perlman (A.C.); the Adelis Foundation(A.C.); and the Irving I. Moskowitz Foundation (A.C.).

## STAR Methods

Contact for reagent and resource sharing

Further information and requests for resources and reagents should be directed to and will be fulfilled by the Lead Contact, Alon Chen.

### Experimental model and subject details

#### Animals

All experiments were approved by and conducted in accordance with the regulations of the local Animal Care and Use Committee (Government of Upper Bavaria, Munich, Germany and Weizmann Institute of Science, Rehovot, Israel).

For all experiments characterizing m^6^A changes after stress, 10–12 w old adult C57 BL/6 male mice were used (Charles River, Sulzfeld, Germany). Mettl3-cKO and Fto-cKO mice were generated by breeding Mettl3^tm1 C(KOMP)Wtsi^ lox/lox mice (Geula et al., 2015) and Fto^tm1 C(EUCOMM)Wtsi^ lox/lox mice obtained from EMMA (EM:05094) to Nex-CreERT2 mice (Agarwal et al., 2012), respectively. Experimental mice were homozygous floxed Nex-CreERT2-positive and Nex-CreERT2-negative littermates generated by breeding of homozygous floxed mice negative and hemizygous for the CreERT2-allele. All experimental animals of the 2 knockout lines were fed with tamoxifen-containing chow (Genobios LASCR diet Cre Active TAM 400) starting at the age of 4–6 w. Animals were housed in groups until being single housed 7 d before the experiments started in standard plastic cages and maintained in a temperature-controlled environment (21 ± 2°C) on a 12 h light/dark cycle with food and water available *ad libitum*. Restraint stress was performed for 15 min in ventilated 50 ml falcon tubes, starting at 2 h post lights on. For pharmacological studies, mice were injected with vehicle solution (saline), 250 μg/kg corticosterone (corticosterone-HBC complex, Sigma) or 10 mg/kg dexamethasone (Ratiopharm Dexa-ratiopharm) i.p. 2 h post switching the lights on.

#### Sample collection

Whole mouse cortex for m^6^A-Seq was collected at designated time points by manual dissection of fresh brains on ice. For each sample, 3 animals randomly selected from the same group were pooled. For investigation of regions-specific effects in PFC and AMY, brains were immediately flash-frozen after dissection and defined tissue punches of medial prefrontal cortex (PFC; consisting of infralimbic and prelimbic cortex) and amygdala (AMY; consisting of central and basolateral amygdala) were collected using a 1 mm round tissue punch while sectioning brains on a cryostat. Mouse whole blood was collected in EDTA tubes, aliquoted and flash-frozen.

#### Cell culture

Human immortalized BLCLs derived from age-matched (33–53 y) male subjects either healthy or diagnosed with MDD were cultured in RPMI-1640 medium (Merck KGaA, Darmstadt, Germany) supplemented with 10 % fetal calf serum at 37 °C with 5 % CO_2_. The cells were tested to be free of mycoplasma. Cells were treated with cortisol (Sigma-Aldrich, St. Louis, MO, in ethanol, final concentration 0.1% v/v) or dexamethasone (Ratiopharm Dexa-ratiopharm, in saline), or ethanol or saline mock control, respectively.

#### Human blood

Human whole blood was collected using PAXgene Blood RNA Tubes (PreAnalytiX, Hombrechtikon, Switzerland) either unstimulated or after oral administration of 1.5 mg dexamethasone and processed as described previously (Menke et al., 2012). Age-matched healthy Caucasian male and females subjects were selected from the “MPIP” and “MARS” cohorts described previously (Arloth et al., 2015; Menke et al., 2012).

### Method Details

#### RNA isolation

Total RNA from tissue, mouse blood and BLCL cells was purified using Trizol (Invitrogen, Life Technologies, Carlsbad, CA) according to the manufacturer’s instructions followed by isopropanol precipitation. For mouse whole blood, RNA was isolated using a 1:10 ratio of blood to Trizol.

#### Global m^6^A measurements

Global m^6^A in total RNA was quantified by the EpiQuik m^6^A RNA Methylation Quantification Kit (Epigentek Group Inc., Farmingdale, NY) following manufacturers' specifications and using 100–300 ng input (in duplicates or triplicates). Comparing total RNA global m^6^A measurements with LC-MS/MS data from the same conditions, we observed high correlation of stress-changes, suggesting that the total RNA colorimetric assay represents an appropriate tool to detect global m^6^A regulation patterns. Brain global methylation in PFC and AMY is not regulated by circadian rhythm (data not shown).

#### LC-MS/MS

Samples were pooled from 4 mice randomly selected from the same group. Residual genomic DNA was removed using the TurboDNA-free kit (Ambion, Life Technologies, Carlsbad, CA). RNA integrity and absence of DNA was confirmed by Bioanalyzer RNA Nano chips (Agilent Technologies, Santa Clara, CA, RIN >9) and Qubit DNA High sensitivity kit (Thermo Fisher Scientific, Waltham, MA), respectively. PolyA+ RNA was prepared using 2 rounds of the Genelute mRNA Prep Kit (Sigma-Aldrich, St. Louis, MO) with rRNA depletion confirmed by Bioanalyzer RNA Nano chips (Agilent Technologies, Santa Clara, CA, mRNA mode). 250 ng PolyA-RNA per sample and a N6-methyladenosine/adenosine standard curve were mixed with deuterated N6-(methyl-d3)-adenosine as an internal spike-in calibrator and processed and measured as reported before (Jia et al., 2011). Quantification was performed by comparison with the standard curve obtained from pure nucleoside standards normalized by the deuterated spike-in calibrator run within the same experiment.

#### m^6^A-Seq

For mouse m^6^A-Seq, whole mouse cortex samples were used pooling 3 individuals each, since m^6^A-Seq on PolyA-RNA of smaller regions did not result in sufficient enrichment quality. For m^6^A-Seq of human BLCLs, RNA from one cell line (healthy male donor) 1 h after treatment with 100 nM cortisol or mock condition was used with 1 technical replicate per condition and no IgG control. m^6^A-Seq was preformed using the previously published m^6^A-Seq-protocol (Dominissini et al., 2013) (BLCL samples) or the following slightly modified version of it (mouse brain): Residual genomic DNA was removed using the TurboDNA-free kit (Ambion, Life Technologies, Carlsbad, CA). RNA integrity and absence of DNA was confirmed by Bioanalyzer RNA Nano chips (Agilent Technologies, St. Louis, MO, RIN > 9.5) and Qubit DNA High sensitivity kit, respectively. PolyA+ RNA was prepared using 1 round of the Genelute mRNA Prep Kit (Sigma-Aldrich, St. Louis, MO) with less than 5 % residual rRNA as confirmed by Bioanalyzer RNA Nano chips (Agilent Technologies, St. Louis, MO, mRNA mode). RNA was fragmented using fragmentation reagent (Life Technologies, Carlsbad, CA). mRNA fragments were precipitated with ethanol and used for m^6^A-immunoprecipitation, IgG control and input samples. m^6^A-immunoprecipitation (10 ug mRNA fragments, 10 ug rabbit polyclonal anti-m^6^A 202 003, Synaptic Systems, Göttingen, Germany) or IgG control (10 ug mRNA fragments mixed from all samples, 10 μg IgG 2729, Cell Signalling Technology, Beverly, MA) was performed in precipitation buffer (50 mM Tris, pH 7.4, 100 mM NaCl, 0.05% NP-40, 1 ml total volume) with 1 μl RNasin Plus (Promega, Madison, WI) rotating head over tail at 4 °C for 2 h, followed by incubation with washed 30 µl Protein A/G beads (Thermo Fisher Scientific, Waltham, MA) rotating at 4 °C for 2 h. Bead-bound antibody-RNA complexes were recovered on a magnetic stand and washed twice with immunoprecipitation buffer, twice with high-salt buffer (50 mM Tris, pH 7.4, 1 M NaCl, 1 mM EDTA, 1 % NP-40, 0.1 % SDS), and twice with immunoprecipitation buffer. Fragments were eluted by Proteinase K treatment (300 μl elution buffer: 5 mM Tris-HCL pH 7.5, 1 mM EDTA pH 8.0, 0.05 % SDS, 4.2 µl 20 mg/ml proteinase K). RNA was recovered from the eluate using Trizol LS (Invitrogen Life Technologies, Carlsbad, CA) following manufacturers' recommendations. Sequencing libraries were prepared using the Illumina TruSeq non-stranded mRNA protocol following the standard protocol starting from mRNA fragments (mouse brain) or the NEBNext Ultra RNA Library Prep Kit for Illumina (NEB, Ipswich, MA). Libraries were quality-checked using Bioanalyzer DNA High Sensitivity chips (Agilent Technologies, St. Louis, MO) and quantified using the KAPA Library Quantification Kit (KAPA Biosystems, Boston, MA). Sequencing was performed on 4 lanes of an Illumina HiSeq4000 PE 2x100 (mouse brain) or on 1 lane of an Illumina MiSeq PE 2x 75 (BLCLs, Illumina, San Diego, CA) multiplexing all m^6^A-, IgG- and input samples.

#### mRNA-Seq

Brains were collected from 5 of each of the following: Mettl3-cKO and WT as well as Fto-cKO and WT mice 24 h after fear conditioning (“FC”, details in “Animal behaviour testing”) or comparable handling without fear induction (“Box”: handling and exposure to context as in “FC” in “Animal behaviour testing” but without foot shock and tone/CS and US). The entire CA1 and CA3 was cryo-punched using 0.7 and 1 mm punching tools from snap-frozen brains sliced at 250 µm using a cryostat and RNA isolated. Residual genomic DNA was removed using the TurboDNA-free kit (Ambion, Life Technologies, Carlsbad, CA). RNA integrity and absence of DNA was confirmed by Bioanalyzer RNA Nano chips (Agilent Technologies, St. Louis, MO, RIN > 8.5) and Qubit DNA High sensitivity kit, respectively. mRNA-Seq libraries were prepared from 4 µg total RNA using the llumina TruSeq stranded mRNA protocol HT (Illumina, San Diego, CA) following the standard protocol starting using Superscript III and 11 cycles of PCR. Libraries were quality-checked using Bioanalyzer DNA High Sensitivity chips (Agilent Technologies, St. Louis, MO) and quantified using the KAPA Library Quantification Kit (KAPA Biosystems, Boston, MA). Sequencing was performed on 4 lanes of an Illumina HiSeq4000 PE 2x100 (Illumina, San Diego, CA) multiplexing all samples.

#### Gene expression

Gene expression of m^6^A-related enzymes was done by SYBR-green-based qPCR. RNA was reverse-transcribed using the SuperScript III VILO cDNA Synthesis Kit (Invitrogen Life Technologies, Carlsbad, CA) and QuantiFast SYBR Green PCR Kit (QIAGEN, Hilden, Germany) on a Quantstudio 7 (Applied Biosystems, Waltham, MA) with the following primers: Mettl3 NM_019721 (ATTGAGAGACTGTCCCCTGG, AGCTTTGTAAGGAAGTGCGT), Mettl14 NM_201638 (AGACGCCTTCATCTCTTTGG, AGCCTCTCGATTTCCTCTGT), Wtap_consensus (GTTATGGCACGGGATGAGTT, ATCTCCTGCTCTTTGGTTGC), Wtap_short NM_001113532 (CTAGCAACCAAAGAGCAGGA, AGTCTTGACTGGGGAGTATGA), Wtap_long NM_001113533 (GGCAAAAAGCTAATGGCGAA, GCTGTCGTGTCTCCTTCAAT), Fto NM_011936 (CTGAGGAAGGAGTGGCATG, TCTCCACCTAAGACTTGTGC), Vir-Kiaa1429 NM_001081183 (CATTACGGCCGCTTAGTTCT, TACCACTGCCTCCACTAACA), Alkbh5 NM_172943 (ACAAGATTAGATGCACCGCG, TGTCCATTTCCAGGATCCGG), Ythdf1 NM_173761 (CATTATGAGAAGCGCCAGGA, AGATGCAACAATCAACCCCG), Ythdf2 NM_145393 (ACCAACTCTAGGGACACTCA, GGATAAGGAGATGCAACCGT), Ythdf3 NM_172677 (TGCACATTATGAAAAGCGTCA, AGATGCGCTGATGAAAACCA), Ythdc1 NM_177680 (TTCATAACATGGGACCACCG, TCATAGTCATGTACTCGTTTATCTC), Hnrnpc NM_016884 (CAAACGTCAGCGTGTTTCAG, TGGGGATGAGAAGGACAAGT), Hnrnpa2 B1 NM_016806 (GTGGAGGGAACTATGGTCCT, TGAAGGCACCAACAAGAACT). Each qPCR assay was performed in duplicates or triplicates with a standard dilution curve of a calibrator and using assay efficiency for calculations. Expression levels were quantified by the ddCT method normalizing to an average of 4–5 housekeeping genes chosen based on maximum stability between conditions from the following: Hprt NM_013556 (ACCTCTCGAAGTGTTGGATACAGG, CTTGCGCTCATCTTAGGCTTTG), Rpl13 A NM_009438 (CACTCTGGAGGAGAAACGGAAGG, GCAGGCATGAGGCAAACAGTC), Atp5j NM_001302213 (TATTGGCCCAGAGTATCAGCA, GGGGTTTGTCGATGACTTCAAAT), Polr2 B NM_153798 (CAAGACAAGGATCATATCTGATGG, AGAGTTTAGACGACGCAGGTG), Rn18s NR_003278 (CAGGATTGACAGATTGATAGC, ATCACAGACCTGTTATTGCTC), Ubc NM_019639 (CTGCCCTCCCACACAAAG, GATGGTCTTACCAGTTAAGGTT), Hmbs NM_001110251 (TCTGAAAGACAGATGGAATGCC, CCACACGGAAAGAGAAGAGGC). For human samples, the following primers were used: NR3C1 NM_000176 (CAGCAGTGAAATGGGCAAAG, TCGTACATGCAGGGTAGAGT), NR3C2 NM_000901 (GATCCAAGTCGTGAAGTGGG, TGAAGGCTGATTTGGTGCAT), FKBP5 NM_004117 (CGGCGACAGGTTCTCTACTT, TCTCCAATCATCGGCGTTTC), TSC22D3 NM_004089 (TCCGTTAAGCTGGACAACAG, TTCAACAGGGTGTTCTCACG) with housekeeping genes TBP NM_003194 (GGGAGCTGTGATGTGAAGTT, GAGCCATTACGTCGTCTTCC), RPL13 A NM_012423 (GCGTCTGAAGCCTACAAGAA, CCTGTTTCCGTAGCCTCATG), and SDHA NM_004168 (CAGGGAAGACTACAAGGTGC, CAGTCAGCCTCGTTCAAAGT).

#### Upstream GRE prediction

10 kb upstream sequences of m^6^A-related genes were retrieved using Biomart (Smedley et al., 2015). GC response elements were predicted by the JASPAR vertebrate core transcription factor binding site prediction (Mathelier et al., 2016) querying NR3C1 motifs MA0113.1 (mammalian), MA0113.2 (mmu), and MA0113.3 (hsa) with a conservative relative profile score threshold of 90 %.

#### Spike-in Oligo

The spike-in RNA oligo was designed with the following specifications: 100 bp length, 3 internal m^6^A sites within GGAC motif flanked by the most frequent nucleotides 5’ U/A, 3’ A/U, not complementary to hsa or mmu RefSeq mRNA or genome, secondary structure exposing m^6^A sites, mean % GC = 51. The sequence is GCAGAACCUAGUAGCGUGUGGmACACGAACAGGUAUCAAUAUGCGGGUAUGG mACUAAAGCAACGUGCGAGAUUACGCUGAGGmACUACAAUCUCAGUUACCA. Fully m^6^A-methylated or unmethylated RNA oligos were purchased from Sigma (Sigma-Aldrich, St. Louis, MO). m^6^A site prediction was performed using SRAMP (Zhou et al., 2016) (full transcript mode, generic predictive model) confirming that the motif sequence context is similar to those occurring in real m^6^A data. Structure prediction was performed using RNAstructure (Reuter and Mathews, 2010) (Fold mode, Version 5.8.1).

#### Candidate m^6^A-RIP-qPCR

To validate m^6^A-Seq experiments, candidates were chosen from the list of differentially methylated transcripts selecting for transcripts with only 1 or few m^6^A-peaks, in the latter case with either only 1 of them stress-regulated or all regulated in a similar manner. For investigation of candidate transcript methylation in small brain areas, candidate lists were constructed by intersecting microarray results of mouse brain PFC, AMY and hippocampus after acute stress and GC stimulation (Arloth et al., 2015) with genes known to be methylated in mouse brain (Hess et al., 2013; Meyer et al., 2012) and functional annotation GO-terms. For investigation of candidate transcript methylation in BLCL cell lines, dexamethasone-responsive genes from human blood microarray data (Arloth et al., 2015) were intersected with BLCL m^6^A-Seq data (unpublished). 15 μg PolyA-RNA (m^6^A-Seq validation) or 3 μg total RNA (candidate m^6^A-RIP-qPCR validation) or 1.5 μg total RNA (brain area/cell line candidate m^6^A-RIP-qPCR) was mixed well with 30 fmol or indicated amount of spike-in or 3 fmol spike-in, respectively, and equally split into 3 conditions: m^6^A-RIP, IgG control and input. For m^6^A-Seq validation and brain are candidate m^6^A-RIP-qPCR only fully methylated spike-in was used. Input samples were flash-frozen during the course of the experiments. m^6^A-RIP and IgG control samples were incubated in parallel to m^6^A-Seq with 1 ug anti-m^6^A antibody (rabbit polyclonal 202 003, Synaptic Systems, Göttingen, Germany) or 1 ug IgG (rabbit polyclonal IgG 2729, Cell Signalling Technology, Beverly, MA) in immunoprecipitation buffer (0.5 ml total volume) with 1 μl RNasin Plus (Promega, Madison, WI) rotating head over tail at 4 °C for 2 h, followed by incubation with washed 25 μl Dynabeads M-280 (Sheep anti-Rabbit IgG Thermo Fisher Scientific, Waltham, MA, 11203 D) rotating head over tail at 4 °C for 2 h. Bead-bound antibody-RNA complexes were recovered on a magnetic stand and washed twice with immunoprecipitation buffer, twice with high-salt buffer, and twice with immunoprecipitation buffer. RNA was eluted directly into Trizol and input RNA was also taken up in Trizol. RNA from all conditions was purified in parallel using the miRNeasy micro RNA isolation kit (QIAGEN, Hilden, Germany) including a 3-time repeated elution 15 μl H_2_O to ensure the complete elution of all RNA. The entire eluate was transcribed to cDNA using the SuperScript III VILO cDNA Synthesis Kit (Invitrogen, Life Technologies, Carlsbad, CA). Gene expression was quantified using TaqMan Fast Advanced Master Mix (Applied Biosystems, Waltham, MA) on a Quantstudio 7 (Applied Biosystems, Waltham, MA) by the following Taqman gene expression assays: Actb NM_007393 (Mm01205647_g1), Akt1 NM_009652 (Mm01331626_m1), Arc NM_018790 (Mm01204954_g1), Atp1 B1 NM_009721 (Mm00437612_m1), Bsn NM_007567 (Mm00464452_m1), Camk2 A NM_009792 (Mm00437967_m1), Camk2n1 NM_025451 (Mm01718423_s1), Cited1 NM_007709 (Mm01235642_g1), Cnr1 NM_007726 (Mm01212171_s1), Crh NM_205769 (Mm04206019_m1), Crhbp NM_198408 (Mm01283832_m1), Crhr1 NM_007762 (Mm00432670_m1), Ctsb NM_007798 (Mm01310508_g1), Cyfip2 NM_133769 (Mm00460148_m1), Dlg4 NM_007864 (Mm00492193_m1), Dnmt1 NM_001199433 (Mm01151063_m1), Dusp1 NM_013642 (Mm00457274_g1), Egr3 NM_018781 (Mm00516979_m1), Fkbp5 NM_010220 (Mm00487406_m1), Fscn1 NM_007984 (Mm00456046_m1), Fth1 NR_073181 (Mm04336020_g1), Gabbr1 NM_019439 (Mm00444578_m1), Gabbr2 NM_001081141 (Mm01352561_m1), Gadd45g NM_011817 (Mm01352550_g1), Grm1 NM_001114333 (Mm00810219_m1), Grm3 NM_181850 (Mm01316764_m1), Homer1 NM_011982 (Mm00516275_m1), Htra1 NM_019564 (Mm00479887_m1), Mllt11 NM_019914 (Mm00480176_m1), Nlgn2 NM_198862 (Mm01245481_g1), Nodal NM_013611 (Mm00443040_m1), Notumos AK028718 (Mm00845023_s1), Nr3c1 NM_008173 (Mm00433832_m1), Nr4a1 NM_010444 (Mm01300401_m1), Nrcam NM_176930 (Mm00663607_m1), Nrxn1 NM_020252 (Mm03808856_m1), Nrxn2 NM_020253 (Mm01236844_g1), Onecut1 NM_008262 (Mm00839394_m1), P2ry13 NM_028808 (Mm00546978_m1), Plekhg3 NM_153804 (Mm00770086_m1), Plin4 NM_020568 (Mm00491061_m1), Pomc NM_008895 (Mm00435874_m1), Prkcb NM_008855 (Mm00435749_m1), Prkcg NM_011102 (Mm00440861_m1), Pvrl3 NM_021495 (Mm01342993_m1), Rgs4 NM_009062 (Mm00501392_g1), Rhou NM_133955 (Mm00505976_m1), Sgk1 NM_001161850 (Mm00441387_g1), Sgk2 NM_013731 (Mm00449845_m1), Sirt2 NM_022432 (Mm01149204_m1), Spats1 NM_027649 (Mm01270591_m1), Sumo1 NM_009460 (Mm01609844_g1), Syn1 NM_013680 (Mm00449772_m1), Syngap1 NM_001281491 (Mm01306145_m1), Tec NM_013689 (Mm00443230_m1), Tsc22d3 NM_001077364 (Mm00726417_s1). Mouse housekeeping genes: Hprt1 NM_013556 (Mm03024075_m1), Rpl13a NM_009438 (Mm01612987_g1), Tbp NM_013684 (Mm01277045_m1), Ubc NM_011664 (Mm02525934_g1), Uchl1 NM_011670 (Mm00495900_m1). Human gene expression assays: ID3 NM_002167 (Hs00171409_m1), DUSP1 NM_004417 (Hs00610256_g1), DDIT4 NM_019058 (Hs01111686_g1), GPER NM_001505 (Hs01922715_s1), IRS2 NM_003749 (Hs00275843_s1), FKBP5 NM_004117 (Hs01561006_m1), NR3C1 NM_000176 (Hs00353740_m1), TSC22D3 NM_004089 (Hs00608272_m1). Human housekeeping genes: RPL13A NM_012423 (Hs04194366_g1), TBP NM_003194 (Hs00427620_m1). The spike-in was quantified using a custom Taqman expression assay (primers TCAATATGCGGGTATGGACTAAAGC, TGAGGACTACAATCTCAGTTACCA and probe AACGTGCGAGATTACG).

#### Human microarray data

Gene expression of m^6^A-related genes was extracted from microarray expression data of human whole blood published previously (Arloth et al., 2015).

#### Animal behaviour testing

All behavioural assessments were performed during the light phase. The experimenter was blinded to the genotype of the animals. Retesting followed the order of least-to-most stressful with 2–3 days’ rest in between tests.

Anxiety-like behaviour was assessed using the Open Field Test (OF, 10 min, 10 lux, grey plastic box 50 × 50 × 50 cm, centre defined as the inner 25 × 25 cm area), Elevated Plus Maze (EPM 5 min, 10 lux on closed arms, 100 lux on open arms, grey plastic maze 50 × 50 cm elevated 25 cm above the floor), Dark Light Box-Test at baseline (DLB basal) and 4 h post 15 min restraint stress (DLB 4 h post stress) (5 min, 100 lux in lit compartment), each with automated tracking (ANY-maze, Stoelting Co., Wood Dale, IL). The Marble Burying Test was performed by placing the mice in a fresh cage with 5 cm flattened fresh bedding with 15 black, clean marbles spaced evenly across (20 min, 10 lux, counting the number of buried marbles every 5 min). Cognitive function was assessed using the Y maze alternation task for working memory (5 min, 10 lux, Y-shaped 3-arm apparatus with 25 cm arm length and distinguishing visual cues on the walls and at the end of each arm, with automated tracking). The proportion of spontaneous non-repeated subsequent entries into each of the 3 arms (alternations) from the total number of 3-arm entries (including repeat entries) was used as the readout. Non-fear-related memory was assessed using the Object-Recognition-Task (ORT, 2x 5 min with 1 h intertrial interval, 10 lux, grey plastic box 50 × 50 × 50 cm, training trial: 2 identical objects with 1 out of 2 objects without object preference randomly assigned to all mice, test trial: 1 known, 1 novel object). The object discrimination ratio DI was determined by DI = (Time with novel object–Time with familiar object) / (Time with novel object + Time with familiar object) within the test trial.

Fear-related memory was assessed by conditional fear learning. Mice were fear conditioned (FC) within the same session for both contextual and cued fear by 180 s of baseline exposure to context A (a metallic/plastic cubic chamber with metal grid conditioned with 70% ethanol smell), followed by a 20 s 80 dB tone (9 kHz sine-wave, conditioned stimulus, CS), which co-terminated with an electric foot shock (unconditioned stimulus, US, 0.7mA, 2s, constant current delivered through the metal grid) and a 60 s after-shock interval. Memory was assessed by measuring freezing in response to the different cues by a highly experienced observer blind to the genotype. Auditory cued fear memory was tested 1 day after FC in context B (cylindrical plastic chamber with bedding conditioned with 1 % acetic acid) by presenting a 3 min CS after 180 s baseline recording and followed by a 60 s post-tone recording. Freezing across the 180 s tone exposure was binned in 60 s intervals to assess short-term stimulus-habituation. Context memory was tested 2 days after FC in context A without presenting US or CS. Fear extinction was achieved by 10*20 s CS presentations (variable inter-trial-interval of 20–60s) in context B on 3 consecutive days 2 w after FC with freezing assessed across the first 3 tone presentations. Fear extinction memory retention was measured 1 w after the extinction by presenting 3*20 s CS and measuring the freezing across those 3 presentations. Animals with generalized fear response (over 50% freezing in any of the baseline recordings of the extinctions trials) were excluded from the analysis for extinction memory.

#### Electrophysiology

Recordings were conducted blind to the animal genotype. Preparation of dorsal hippocampal slices and electrophysiological measurements were performed according to standard procedures as we described previously (Schmidt et al., 2011). From every animal, 2 slices were used for the experiments.

#### *In situ* hybridization

Expression quantification of Mettl3 mRNA and Fto mRNA in Mettl3-cKO and Fto-cKO animals was performed by *in situ* hybridization using S-35 labelled antisense probes targeting the floxed exon as described previously (Refojo et al., 2011). Probes were designed for Mettl3 NM_019721 exon 4 (probe cloned using TCAGTCAGGAGATCCTAGAGCTATT and CTGAAGTGCAGCTTGCGACA) and Fto NM_011936 exon 4 (probe cloned using TGGCAGCTGAAATACCCTAAACT and ATAGCTGTACACTGCCACGG). Slides were exposed to Kodak Biomax MR films (Eastman Kodak Co., Rochester, NY), developed, and autoradiographs digitized and quantified by optical densitometry of 2 slides each averaging the signal across both hemispheres and slides utilizing ImageJ (dorsal: Bregma −1.82, −1.94; ventral Bregma −3.16, −3.28).

#### Western Blot

Cells were lysed on ice in RIPA buffer (150 mM NaCl, 1 % NP-40, 0,5 % Sodium deoxycholate, 0,1 % SDS, 50 mM Tris-HCl pH 8 with cOmplete, EDTA-free Protease Inhibitor Cocktail Mini, Roche Applied Science, Roche Diagnotics, Indianapolis, IN) for 30 min. 25 μg total protein as determined by the Bio-Rad Quick Start Bradford Kit (Bio-Rad Laboratories Inc., Hercules CA) was heated for 10 min in SDS/PAGE sample buffer (final concentration 62.5 mM Tris-HCl pH 6.8, 2% SDS, 10% glycerol, 5% b-mercapto-ethanol), separated on a Tris-Glycine SDS–PAGE (Bio-Rad Laboratories Inc., Hercules, CA) and transferred to a nitrocellulose membrane (Amersham Protran, Millipore, Billerica, MA). Membranes were blocked for 1 h in TBST containing 5% non-fat milk, followed by incubation with primary antibodies overnight at 4 °C (anti-GR monoclonal rabbit, ab109022, 1:50000; anti-BTUB polyclonal rabbit, ab6046, 1:10000; Abcam, Cambridge, UK) in TBST with 3% non-fat milk. After incubation with horseradish-peroxidase-coupled secondary antibody (Cell Signalling Technology, Beverly, MA; 7074) at room temperature for 2 h, immunoblots were visualized using enhanced chemiluminescence (ECL Plus, GE Healthcare Life Sciences, Freiburg, Germany). Band intensity was quantified using ImageJ.

## Quantification and statistical analysis

### m^6^A-Seq analysis

Sequencing data quality control was performed by FASTQC (Andrews, 2010). Genomic alignment was performed for the mouse brain samples using the STAR aligner (Dobin et al., 2013) (to Gencode M11/Ensemble 86 build, mm10, at default settings using only those reads mapping uniquely to the genome and with a Phred quality score R 20) and for BLCL samples using TopHat2 (Kim et al., 2013) (to UCSC GRCh37*/*hg19 at default settings). We used exomePeak (Meng et al., 2014) for peak calling for both stress and basal condition or cortisol and mock condition, computing the enrichment of m^6^A-Seq signal in relation to the baseline RNA-Seq (input) signal at a given position, for mouse samples using high-confidence peaks called across all biological replicates and working in transcriptome space. Peaks were annotated using ChIPpeakAnno (Zhu et al., 2010). Upon inspection, condition-unique peaks had some enrichment in the m^6^A-Seq in both stress- and basal conditions arguing for condition-unique peaks being rather caused by peak-detection thresholds than being true present/absent peaks. Therefore, we continued our analyses merging both peak sets (GRanges (Lawrence et al., 2013)). Differential m^6^A expression was analysed using DRME (Liu et al., 2016), a tool specifically developed to assess methylation changes on RNA. IgG control libraries had a low percentage of alignment with no specific enrichments present. Differential gene expression was evaluated using the EdgeR package (Robinson et al., 2010). Calculations and plots were done using R (R Development Core Team, 2011) and ggplot2 (Wickham, 2009). For mouse samples, only genes with an average estimated RPKM>1 in the input samples (Reads Per Kilobase of transcript per Million mapped reads; estimated by EdgeR rpkm function using the longest annotated isoform from mm10) were considered for both DGE (13,090 of 16,031 detected expressed genes) and m^6^A-peak calling (25,821 mapping to 11,534 genes of unfiltered 28,725 detected peaks mapping to 13,536 genes). Both peaks and genes expressed were considered differential with an absolute fold change > 0.1 and a Benjamini-Hochberg corrected P-value=Q < 0.05 for mouse samples and fold change > 1 and a Benjamini-Hochberg corrected P-value=Q < 0.05 for human samples. Distribution-plots of m^6^A across the transcript length were evaluated using the Guitar plots (Cui et al., 2016) package. Our data revealed a higher peak abundancy within the 5’UTR than previously reported which may be due to higher representation of full length 5’UTRs in our input RNA compared to earlier studies (Meyer et al., 2012) and a high contribution of m^6^Am peaks (Linder et al., 2015). GO-term overrepresentation was calculated using the PANTHER Overrepresentation Test (Mi et al., 2013) for “GO biological process complete” and “PANTHER Pathways” with the list of all detected genes as background. Motif search was performed by DREME (Bailey, 2011) and CentriMo (Bailey and Machanick, 2012) using all detected m^6^A peaks as input (mouse: 200nt sequences centred on peak summit). Motif across the whole manuscript are presented with R=A/G, W=A/T, K=G/T, B=G/C/T, **H**=A/C/T, D=A/G/T, V=A/G/C. For comparison of the detected mouse m^6^A-Seq GGACWB motif with known motifs, we employed Tomtom (Gupta et al., 2007; Ray et al., 2013) and CentriMo (Bailey and Machanick, 2012). Comparison of peaks to known m^6^A was done using m^6^A data from RMBase (Sun et al., 2016) (data set of 2015-10-20, GSE29714 mouse brain (Meyer et al., 2012), GSE71154 mouse brain and liver (Ke et al., 2015), SRA280261 mouse embryonic fibroblasts (Zhou et al., 2015), GSE63753 mouse liver (Linder et al., 2015)) using GRanges (Lawrence et al., 2013). Peak position was annotated with Biomart (Smedley et al., 2015) and ChIPpeakAnno (Zhu et al., 2010). Peaks on 5’UTR-CDS border and CDS-3’UTR border are counted as 5’UTR and 3’UTR, respectively. Sequencing tracks were visualized with the UCSC browser (Kent et al., 2002). Overlap with existing m^6^A-reader and FMRP/FMR1 PAR-CLIP and HITS-CLIP data was done using data sets for: YTHDF1 (Wang et al., 2015), YTHDF2 (Wang et al., 2014), YTHDF3 (Shi et al., 2017), YTHDC1 (Xu et al., 2014), HNRNPC (Liu et al., 2015), HNRNPA2 B1 (Goodarzi et al., 2012), FMRP (Ascano et al., 2012), FMR1 (Darnell et al., 2011). To compare human and mouse m^6^A-peaks and binding sites gene symbols were used, first intersecting all datasets with a background list consisting of only human homologues with genes expressed in our m^6^A-Seq (Smedley et al., 2015) and genes expressed in the cell lines from which the CLIP data is derived from (Higareda-Almaraz et al., 2013; Sultan et al., 2008). For random models, an appropriate number of genes were randomly selected from this background list.

### mRNA-Seq

Sequencing data quality control was performed by FASTQC (Andrews, 2010). Reads were quality filtered (Q 20) and adapter trimmed using Cutadapt (Martin, 2011) and pseudoaligned to the mouse transcriptome (Gencode M15 transcripts GRCm38.p5) using kallisto (Bray et al., 2016; using an index with k-mer length 31, paired end strand-specific alignment, sequence based bias correction and 100 bootstraps). Differential gene expression analysis was performed using DESeq2 (Love et al., 2014) after importing the counts and summarizing to gene level with tximport (Soneson et al., 2015; considering genes with a minimum of 25 raw counts in all samples only). ENSMUSG00000019768 and ENSMUSG00000037984 were excluded from the differential gene expression analysis. For analysis of basal expression patterns, as presented in Figure 4, only the subset of unstressed “Box” animals was used. Genes expressed were considered differential with an absolute fold change > 0.2 and a Benjamini-Hochberg corrected P-value=Q < 0.1.

### Gene expression

Statistics were performed on log2 normalized data using a 2x2 MANOVA in SPSS and were multiple testing-corrected by the Benjamini-Hochberg test (cut-off Q<0.05) in R and a cut-off by effect size (η²> 0.01) and post-hoc testing (Tukey HSD).

### Candidate m^6^A-RIP-qPCR

RNA abundance levels were quantified from the input samples using the ddCT method normalizing to the average of all housekeeping genes. Immunoprecipitation efficiency for each biological sample was assessed using the measured abundance of spike-in per m^6^A-RIP/IgG control sample. Because all conditions per sample were equally split at the beginning, % methylation or IgG signal was calculated as follows:

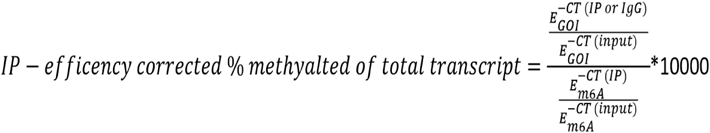

Statistics were performed on log2 normalized data (RNA) or absolute values (m^6^A) using a 2x2 MANOVA in SPSS with multiple testing performed using the Benjamini-Hochberg method (cut-off Q<0.05) in R and a cut-off by effect size (η²> 0.01) and post-hoc testing (Tukey HSD).

## Statistical analysis

Statistical tests were performed using SPSS (IBM SPSS Statistics, Armonk, NY: IBM Corp.) and R (R Development Core Team, 2011) as indicated in Figure legends with n and statistical results indicated in Figure legends und Supplemental Tables. Plots were produced with R (R Development Core Team, 2011) ggplot2 (Wickham, 2009) with definition of presented measurements indicated in the Figure legends. For animal experiments, sample size was estimated a priori using G*Power (Faul et al., 2007) using α = 0.05 and an experience-based β. Animals and samples within experiments were randomized using stratified randomization assisted by random number generation.

## Data availability

All supporting data for this study are available from the corresponding author upon request. Sequencing data will be deposited at GEO repositories before publication.

## Supplementary Information

### Supplementary Tables

Table S1: m^6^A peaks. Related to Figures 1 and S1.

Table S2: Statistics (GLMs & ANOVAs). Related to Figures 1-3 and S1-3.

Table S3: mRNA-Seq of Mettl3-cKO and Fto-cKO mice. Related to Figures 4-5 and S4-5.

### Supplementary Figures

**Figure S1.**
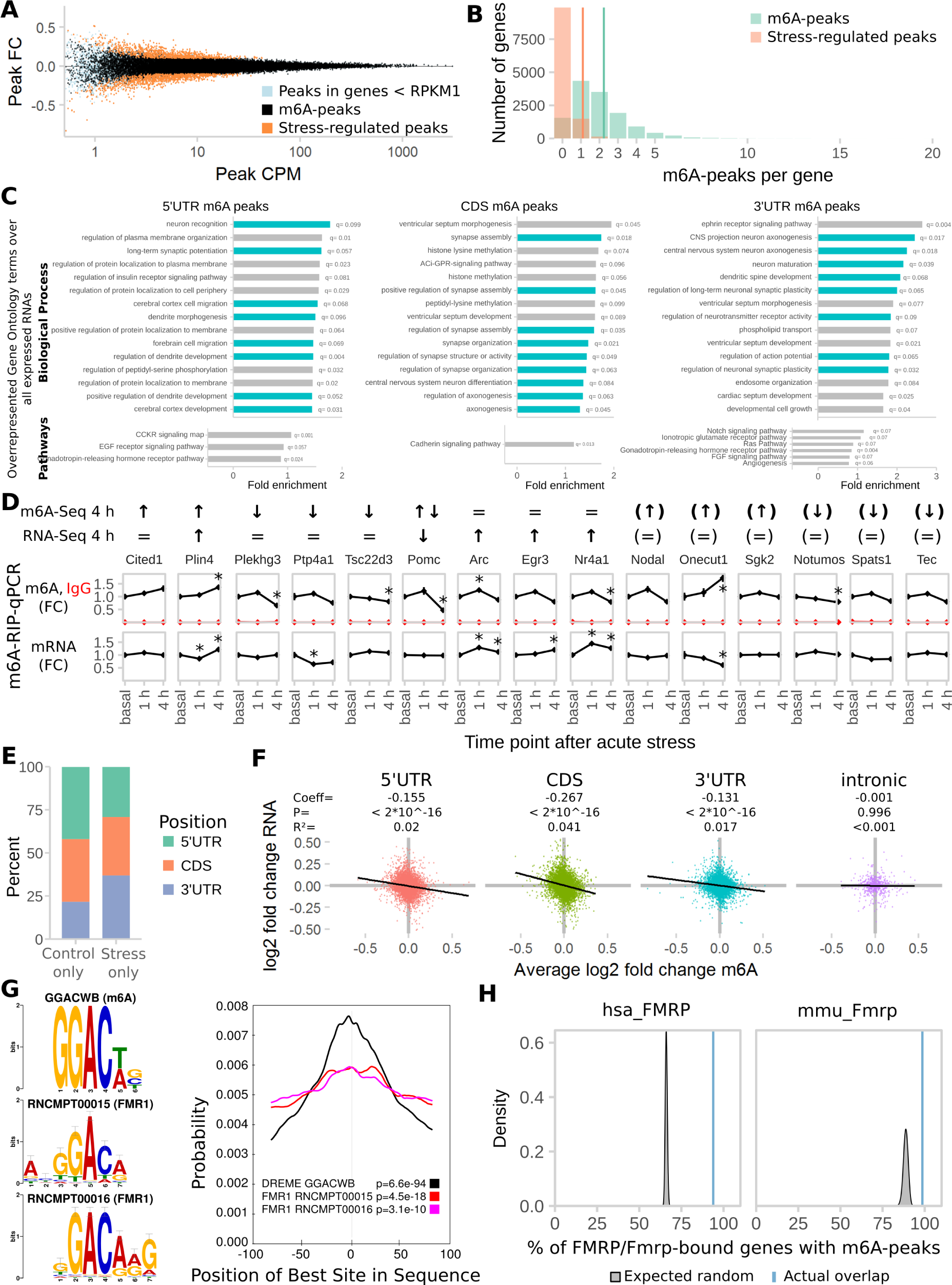
Additional analysis of the M6A-Seq data. Related to Figure 1. **(A) MA Plot of detected m^6^A-peaks showing the distribution of regulated m^6^A-peaks.** Peaks in genes with RNA abundancy of Reads Per Kilobase of transcript per Million mapped reads (RPKM)<1 were removed before analysis (light blue). FC= fold change, CPM=counts per million. **(B) Distribution of number per m^6^A-peaks per transcript.** Methylated genes had 1–20 peaks per gene with average = 2.24. In contrast, on genes with stress-regulated m^6^A, on average 1.1 peaks per transcript were regulated. **(C) Stress-regulated m^6^A-peaks are highly enriched in genes connected to neuronal development, neuronal plasticity, and different cellular pathways with distinct functions depending on the peak position.** (15 highest enriched Biological Process and Pathway gene ontology (GO) terms. Synaptic-plasticity-related GO-terms marked in blue. Overrepresentation test of m^6^A-peaks by position test compared to all genes detected in input samples with FDR-corrected Q<0.1). **(D) Full set of differential m^6^A-peaks chosen for validation with m^6^A-RIP-qPCR showing that many differences found by m^6^A-Seq can be recapitulated on full-length transcript methylation.** m^6^A-Seq 4 h indicates whether and in which direction any of the genes m^6^A-peaks were found to be regulated. RNA-Seq 4 h indicates regulation of the gene mRNA. Arrows in brackets indicate genes found to be regulated but expressed below the cut-off of RNA abundancy RPKM>1. (n = 7, absolute FC>0.1, Q<0.05 with FDR-correction of peaks to respective total peak-set with or without RPKM filter). m^6^A-RIP-qPCR panels show the measured full-length m^6^A-levels and RNA-levels of the respective genes. To illustrate the time-dependency of m^6^A-regulation by stress we also measured transcript methylation 1 h after stress in m^6^A-RIP-qPCR. (Separate cohort of mice, n = 7, mean ± SEM, * depict omnibus Tukey post-hoc tests to basal P<0.05 after FDR-corrected one-way ANOVA. Con = Control, Str = Stress). **(E) Control-specific peaks are enriched in 5’UTR position whereas stress-specific peaks are enriched in 3’UTR position.** **(F) CDS and 5’UTR peaks contribute most to the negative correlation of m^6^A change and RNA change.** (log2 fold changes of m^6^A and RNA after stress. Black line: Linear model + 95% CI. GLMs see Supplementary Table 2). **(F) Comparing the m^6^A-motif GGACWB to known motifs of RNA-binding proteins, 2 motifs for FMR1 are found to be most colocalized.** (Tomtom motif top 2 comparison results: RNCMPT00015 (P = 1.03 E-02, E = 2.52), RNCMPT00016 (P = 1.03 E-02, E = 2.52). Centrimo results: Both motifs are centrally enriched in m^6^A-peaks). **(G) Previously reported genes bound by human and mouse FMRP/FMR1 are enriched for m^6^A-bound genes found in our study.** (The amount of overlap observed (blue line) was compared to distributions gained from 100 random permutations (grey distributions) of all observed expressed genes or all observed expressed genes with human homologues (Z-Test)).

**Figure S2.**
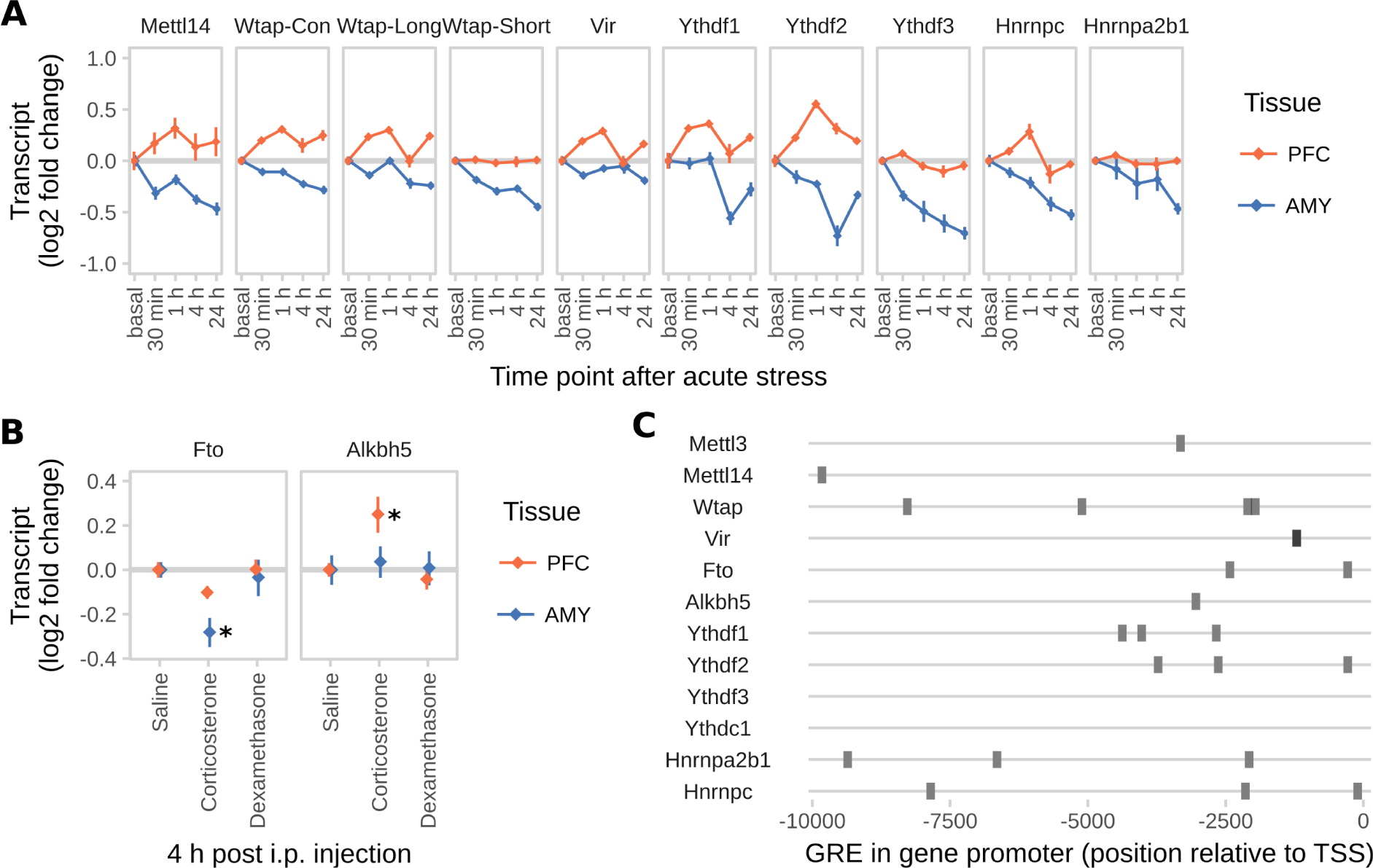
Acute injection of corticosterone i.p. leads to similar changes in global m^6^A in the PFC and AMY like acute stress, suggesting that the effect on m^6^A is mainly mediated by glucocorticoids. Related to Figure 2. **(A) Several m^6^A-related genes are not regulated by acute stress indicating specificity of stress effects.** Wtap expression is measured specifically for the long and short isoform as well as with primers measuring both (Con). (n = 12, log2 fold change ± SEM. 2-way MANOVA without significant interaction or main stress effects (FDR-corrected P< 0.05 and n²>0.01). Full statistics see Supplementary Table 2). **(B) Gene expression regulation of m^6^A-demethylases Fto and Alkbh5 in the PFC and AMY shows similar patterns of regulation after corticosterone injection like after acute stress.** (Fold change measured with qPCR; n = 12, mean ± SEM. Kruskal-Wallis-Test PFC Alkbh5 and AMY Fto P<0.05, Stars: omnibus post-hoc comparisons to basal, P<0.065). **(C) The majority of m^6^A regulatory genes have upstream Glucocorticoid Response Elements (GRE).** Prediction of high confidence GRE sites based on GRE consensus motif MA0113 10 kb upstream of the transcription start site (JASPAR, 90% relative profile score threshold).

**Figure S3.**
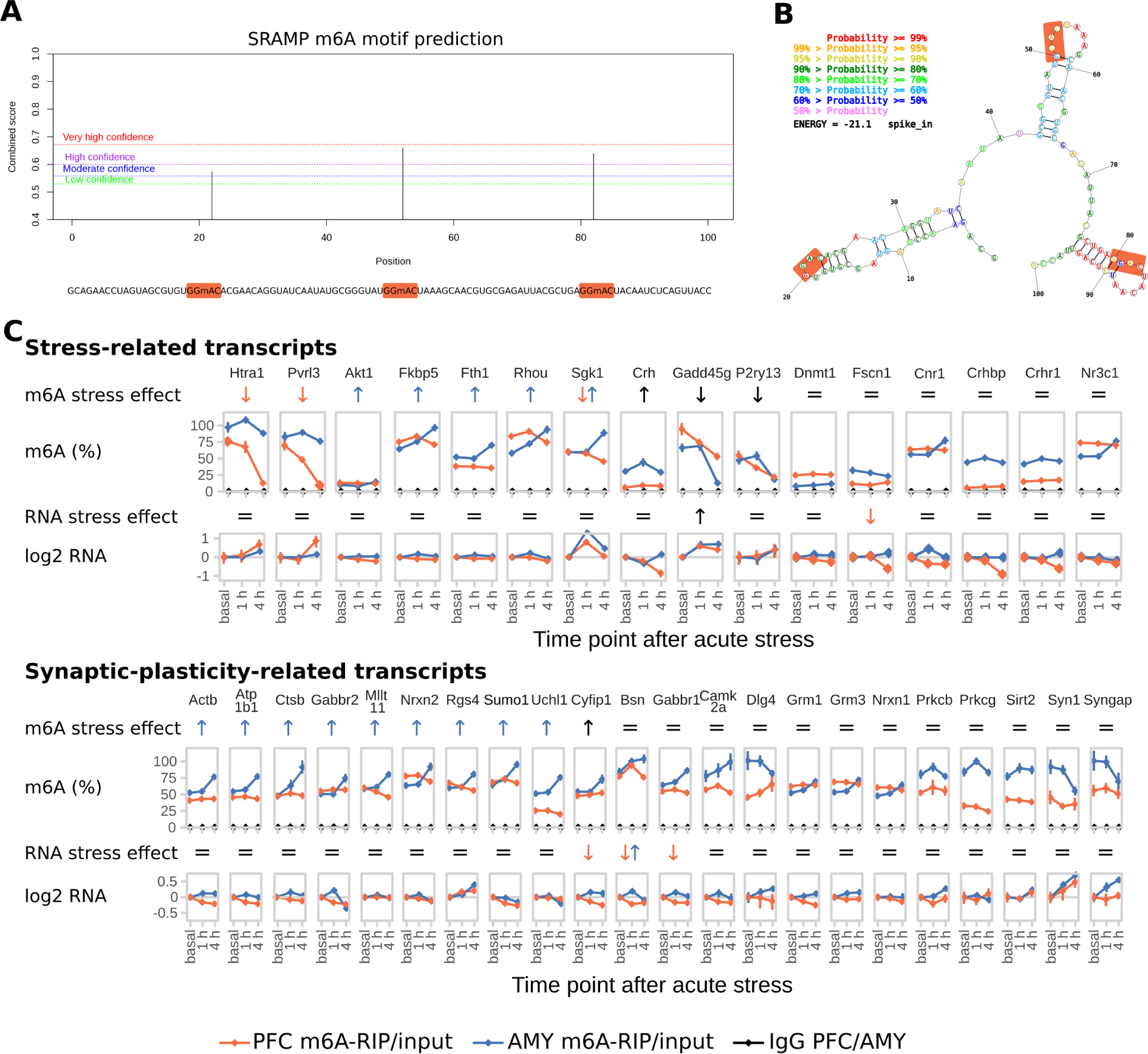
In-depth analysis of the M6A-RIP-qPCR data. Related to Figure 3. **(A) Sequence and m^6^A-site prediction of the synthetic spike-in oligo.** The GGAC consensus motif containing the m^6^A sites is marked up in the sequence string. **(B) Maximum free energy secondary structure** of the oligo. **(C) Absolute full length m^6^A-levels of stress-related and synaptic plasticity-related transcripts are differentially regulated in PFC and AMY of stress-related candidate transcripts and synaptic-plasticity-related candidate transcripts after stress.** Extended data from Figure 3. % m^6^A = % expression after precipitation relative to the total abundancy in input, normalized for immunoprecipitation efficiency by an internal methylated spike-in control. log2 RNA = log2 fold changes of transcript in input samples normalized to 5 housekeeping genes. (n = 8, mean ± SEM. Significant effects observed in FDR-corrected 2-way MANOVA (P<0.05, n²>0.01) are coded in the rows “m^6^A stress effect” and “RNA stress effect”: orange/blue arrows = PFC-/AMY-specific stress effect (interaction effect 2-way ANOVA, one-way follow up significant in respective tissue), black arrow = stress main stress effect, equals sign = no interaction or stress main effect in 2-way ANOVA. For full statistics see Supplementary Table 2).

**Figure S4.**
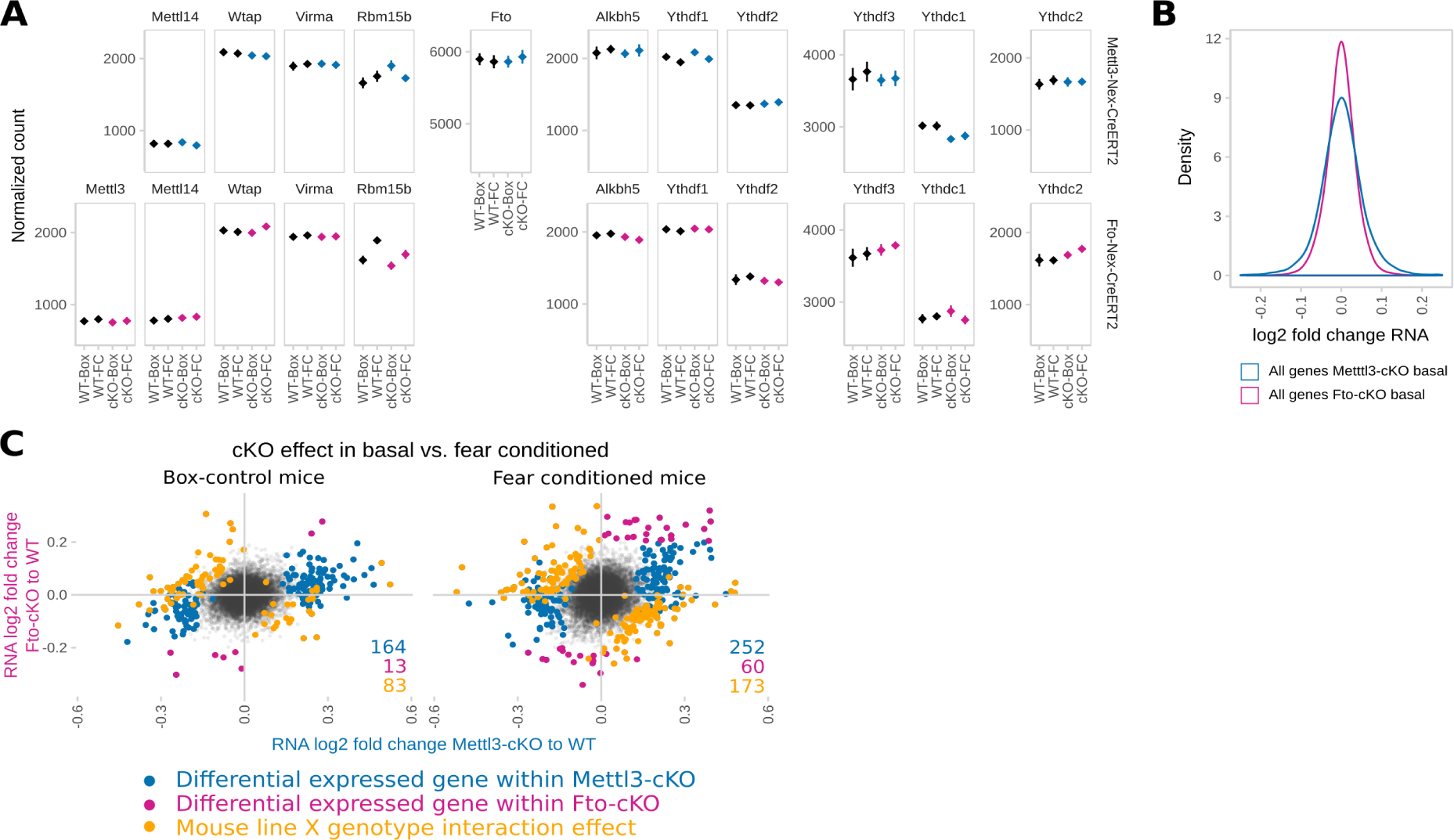
Absence of compensation by other enzymes and further transcriptomic properties in Mettl3- and Fto-cKO mice. **(A) Depletion of *Mettl3* or *Fto* in adult excitatory neurons is not compensated by changes of expression in other genes catalysing and or binding m6A nor is the expression of those genes changed 24 h after fear conditioning.** (DeSeq2 Normalized counts of genes plotted across both Mettl3-cKOs and Fto-cKOs and respective wild type animals (WT) including animals 24h after fear conditioning (FC) and control animals (Box). n=5. Significant genotype contrasts with T-Test P<0.05 before but not after multiple testing correction: *Rbm15 B* in Box-Mettl3-cKOs, *Rbm15 B* in FC-Fto-cKOs, *Ythdc1* in Box MEttl3-cKOs, *Ythdc2* in FC-Fto-cKOs, significant fear conditioning contrast T-Test P<0.05 before but not after multiple testing correction: *Rbm15 B* in WT Fto-cKO and cKO-Mettl3 animals). **(B) Mettl3-cKO gene expression is more variable than Fto-cKO gene expression plotted across all quantified genes.** (Distribution of individual gene expression changes independent of differential calling in Fto-cKOs compared to Mettl3-cKOs. n=5) **(C) Gene expression changes in Mettl3-cKOs compared to their respective gene expression change in Fto-cKOs are more diverse in fear conditioned animals than in unstressed Box-control animals.** (Differentially expressed genes marked by colour: blue = genes differentially expressed in Mettl3-cKOs compared to WT, pink = genes differentially expressed in Fto-cKOs compared to WT, orange= genes expressed in a mouse line x genotype fashion. n=5)

**Figure S5.**
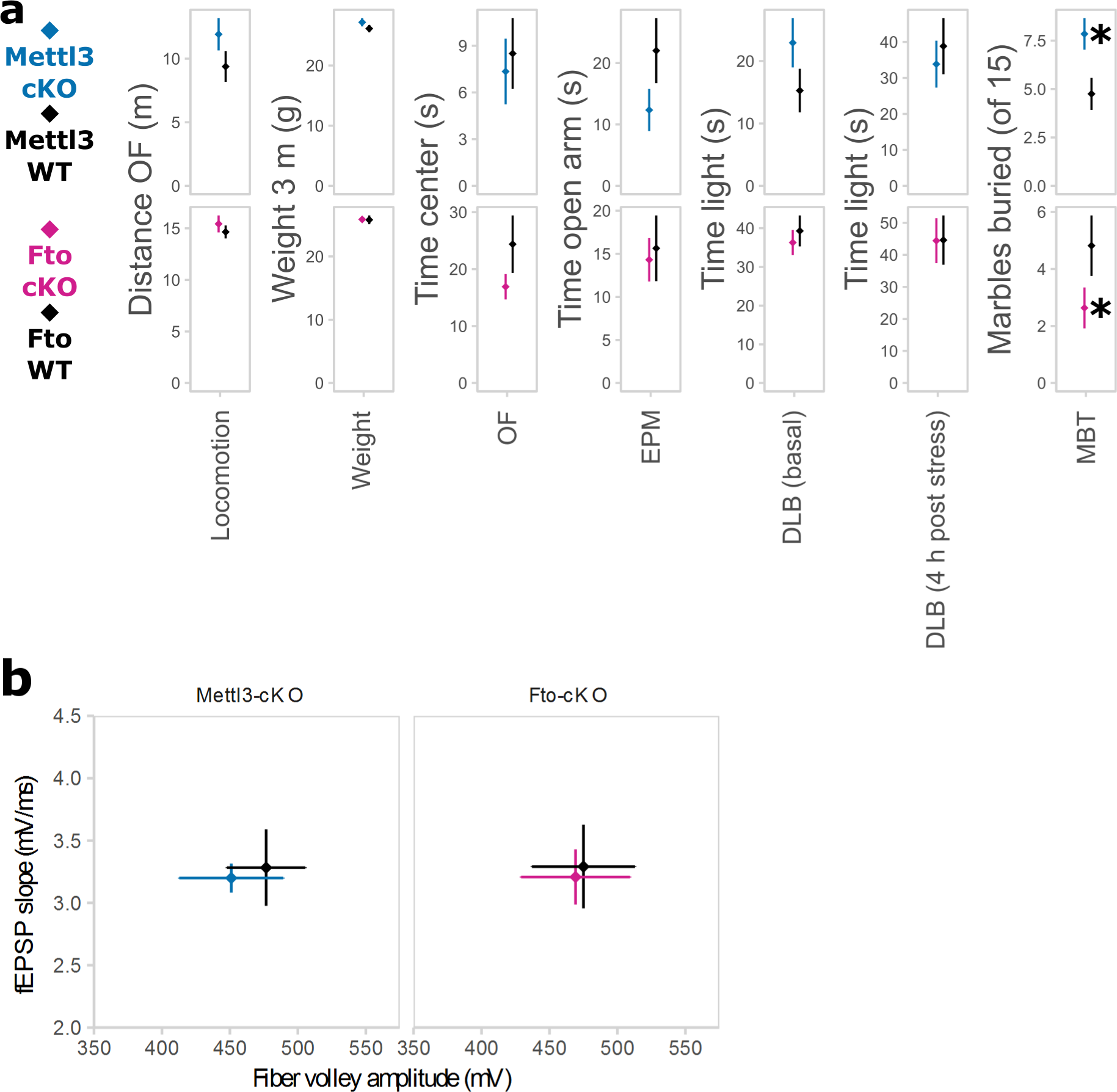
Anxiety-like behaviour is not changed in Mettl3-cKO and Fto-cKO animals. (A) cKO animals did not differ in locomotion, weight or several measurements of anxiety-like behaviour, but spontaneous digging behaviour. OF = Open Field Test, EPM = Elevated Plus Maze, DLB = Dark Light Box, MBT = Marble Burying Test, WT = wild type animals, cKO = conditional knockout animals. Spontaneous burying behaviour as measured by the MBT was increased in Mettl3-cKO animals while decreased in Fto-cKO animals. Weight 6 w post induction with Tamoxifen (average 12 w of age). Marbles buried within 10 min. (n = 11–13, mean ± SEM. * depict T-Tests P<0.05).

**Figure S6.**
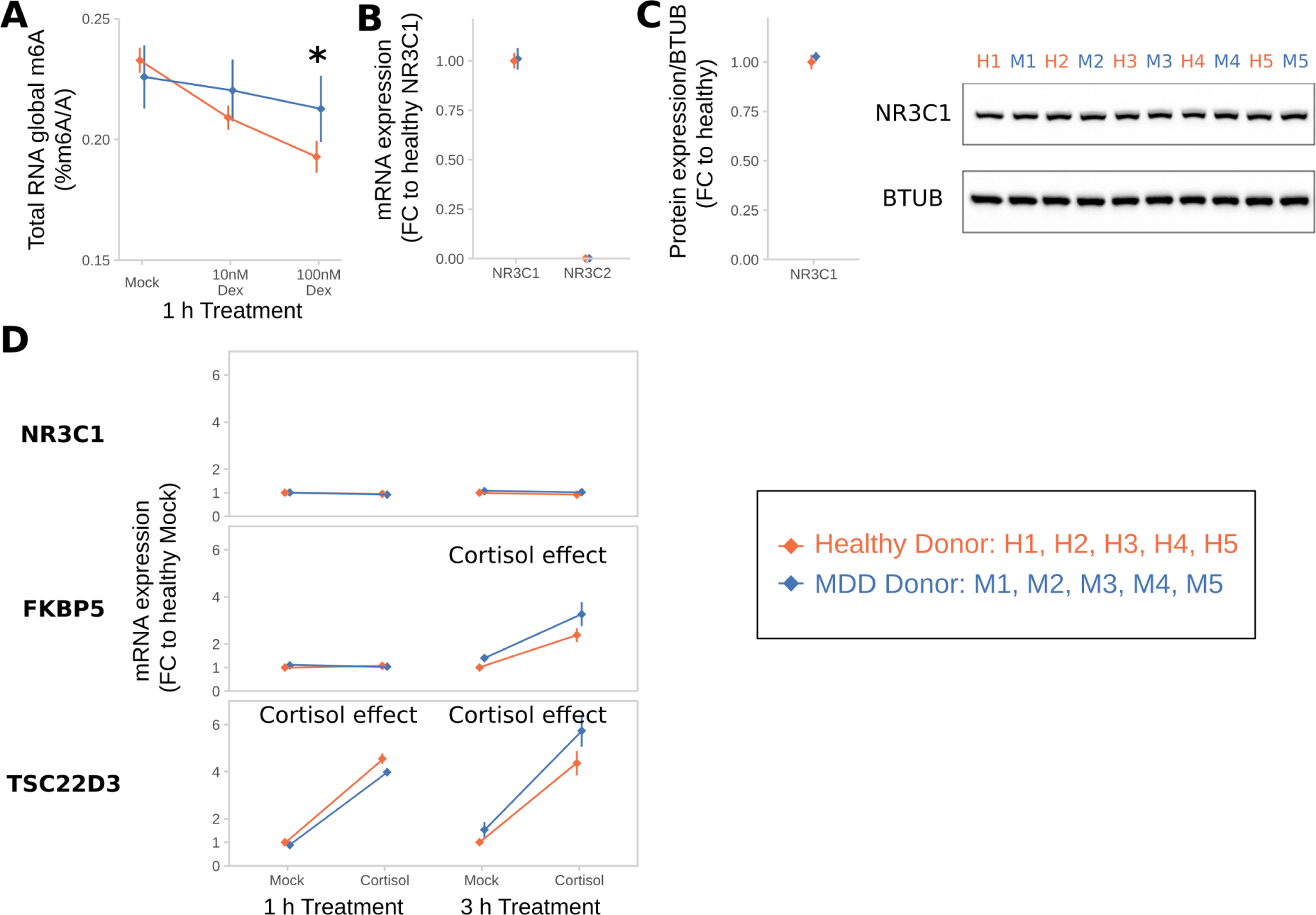
B lymphocyte cell lines (BLCLs) of Major Depressive Disorder (MDD)-donors do not downregulate m^6^A upon Dexamethasone-stimulation. MDD-BLCLs have normal NR3C1-levels and glucocorticoid responsivity. Related to Figure 6. Dex = dexamethasone. **(A) Global m^6^A in BLCLs after dexamethasone treatment is decreased in BLCLs from healthy, but not MDD-donors.** (Global m^6^A assay on total RNA, n = 5 biological replicates with 3 technical replicates each, mean ± SEM. 2-way ANOVA: significant interaction effect of Dex and donor status (F(3,24) = 10.127, P=0.001). * depicts omnibus Tukey post-hoc tests to basal P<0.05). **(B) BLCLs from healthy and MDD donors have comparable levels of NR3C1 mRNA. Levels of NR3C2 are very low but also unchanged.** (qPCR, n = 5 biological replicates, mean ± SEM). **(C) BLCLs from healthy and MDD donors have comparable levels of NR3C1 protein.** (Western Blot quantification of NR3C1 relative to B-TUBULIN (BTUB), n = 5 biological replicates, mean ± SEM). **(D) BLCLs from healthy and MDD donors upregulate FKBP5 and TSC22D3 after cortisol-treatment (100 nM) in the same way.** (qPCR, n = 5 biological replicates, mean ± SEM. 2-way ANOVA: “Cortisol effect” indicates a significant main effect of cortisol treatment: FKBP5 3 H: F(1,16)=13.171, P<0.001, TSC22D3 1 H: F(1,16)=55.245, P<0.001, TSC22D3 3 H: F(1,16)=71.518, P<0.001).

